# Biuret inhibits Arabidopsis root growth through an active, reversible, and genetically tractable developmental response

**DOI:** 10.64898/2026.06.29.735191

**Authors:** Virginia Protto, Virginie Thiry, Arnaud Didier, Thibaut Perez, Gabriel Krouk, Benoit Lacombe, Anna Medici

## Abstract

Biuret, a nitrogen-rich by-product of urea and a common contaminant of urea-based fertilisers, has long been considered a passive phytotoxin, affecting plant performances. Yet its effects on root development and the existence of endogenous mechanisms of perception or tolerance remain largely uncharacterised. Here we combined physiological, developmental, genetic and transcriptomic approaches to investigate the response of *Arabidopsis thaliana* to biuret. Biuret inhibited primary root growth in a dose-dependent manner by reducing meristematic cell division rather than cell elongation, and concomitantly impaired shoot growth by limiting leaf expansion. This root inhibition was reversible upon biuret removal and was accompanied by increased auxin-responsive (*DR5*) and decreased cytokinin-responsive (*TCS*) outputs at the root apex, consistent with a regulated remodelling of meristem activity rather than purely cumulative damage. A forward genetic screen identified the biuret-resistant mutant *bir29*, which sustained root and inflorescence development under inhibitory concentrations. Using ¹⁵N-labelled biuret, we showed that resistance occurred without any change in biuret influx or accumulation, uncoupling sensitivity from exposure. Whole-genome transcriptomics revealed that *bir29* fails to execute the wild-type response, neither repressing the cell-cycle machinery nor deploying the stress-associated programme induced by biuret. Genetic characterisation linked resistance to multiple genomic loci required for full resistance. Together, the results indicate that biuret triggers an active, reversible and genetically tractable developmental response, suggesting that this xenobiotic compound is integrated into endogenous signalling networks.

**Significance Statement:** Biuret, a poorly metabolised contaminant of urea fertilisers, is generally regarded as a passive phytotoxin, yet we show that it inhibits *Arabidopsis* root growth through a reversible and genetically tractable developmental response, accompanied by reorganised auxin and cytokinin signalling, rather than through cumulative chemical injury. The isolation of the resistant mutant *bir29* suggests that plants integrate this xenobiotic molecule into endogenous signalling networks, reframing biuret as an informative probe of root developmental regulation.

## Introduction

Biuret (carbamoylurea, C₂H₅N₃O₂) is a nitrogen-rich, largely anthropogenic compound formed by the condensation of two urea molecules during the industrial synthesis and granulation of urea, of which it is a common impurity and the principal route into agroecosystems (Mikkelsen, 1990; Ochiai *et al*., 2020). Biuret can also arise in soils from the microbial catabolism of cyanuric acid and s-triazine herbicides, and many microorganisms hydrolyse and use it as a nitrogen source (Aukema *et al*., 2020; Robinson *et al*., 2018; Seffernick and Wackett, 2016). Plants are thus exposed to biuret both as a fertilizer-associated compound and as a product of soil microbial activity (Aukema *et al*., 2020; Mikkelsen, 1990; Ochiai *et al*., 2020), and it can induce injury where tissues encounter locally elevated concentrations, for example during direct seeding or foliar urea sprays (Ferreira *et al*., 1986; Mikkelsen, 1990; Reeves *et al*., 1977). Although modern formulations keep biuret low (≤1.2% in urea; ≤0.25% for foliar use on sensitive crops) and thereby limit severe agronomic toxicity, plant responses to biuret remain poorly characterized, particularly at the developmental and molecular levels (Mikkelsen, 1990).

To date, the effects of biuret have been characterized mainly in shoots, in seeds during germination and seedling establishment, and their severity varies markedly between species (Mikkelsen, 1990). Typical symptoms include leaf chlorosis, “yellow-tip” syndrome and, in direct-seeded rice, severe albinism linked to disrupted chloroplast development and impaired photosynthesis (Achor and Albrigo, 2005; Mikkelsen, 1990; Zhang *et al*., 2023). Early seedling growth appears particularly sensitive to biuret exposure, suggesting that actively developing tissues are especially vulnerable to this compound (Ochiai *et al*., 2020; Wilkinson and Ohlrogge, 1960). Biuret accumulation has long been associated with perturbations of nitrogen metabolism and, in some species, with altered protein synthesis (Mikkelsen, 1990); accordingly, rice exposed to excess biuret accumulates the nitrogen-rich compounds allantoin and citrulline, a response thought to facilitate the assimilation and detoxification of excess ammonium through its incorporation into organic molecules (Ochiai *et al*., 2022). A likely explanation for this toxicity is that, unlike many soil microorganisms and some fungi, archaea and green algae, no dedicated biuret hydrolase has so far been identified in land plants and therefore cannot efficiently catabolize it (Ochiai *et al*., 2020; Robinson *et al*., 2018). Consistent with this, heterologous expression of a bacterial biuret hydrolase is sufficient to confer tolerance and to address biuret-derived nitrogen into plant metabolism in rice, indicating that toxicity reflects biuret accumulation within plant tissues rather than a purely external, rhizospheric effect (Ochiai *et al*., 2020). How biuret affects root growth, and whether plants possess endogenous mechanisms to perceive or tolerate it, nevertheless remain unresolved.

Primary root growth is sustained by the root apical meristem (RAM), in which a stem-cell niche organized around the quiescent centre (QC) continuously generates new cells. Root growth reflects the balance between cell division in the proximal meristem and subsequent cell expansion, while meristem size is set by the relative rates of division and differentiation (Blilou *et al*., 2005; Motte *et al*., 2019). Because the RAM integrates developmental and environmental information into growth decisions, it offers a highly sensitive readout of how chemical perturbations affect cell proliferation and meristem activity (Colón-Carmona *et al*., 1999; Motte *et al*., 2019).

Among the major regulators of RAM organization, auxin and cytokinin act centrally and antagonistically (Overvoorde *et al*., 2010; Wybouw and De Rybel, 2019). A local auxin maximum at the root tip maintains stem-cell identity and sustains division, whereas cytokinin drives the transition from proliferation to differentiation at the proximal meristem boundary; the spatial balance between the two therefore defines meristem size and shapes root growth dynamics (Benková *et al*., 2003; Blilou *et al*., 2005; Wybouw and De Rybel, 2019; Zürcher *et al*., 2013). The auxin–cytokinin balance thus provides a useful framework for interpreting altered RAM activity under stress.

Roots, however, do not respond to environmental constraints passively. They continuously sense abiotic and biotic cues and integrate them into adaptive developmental and metabolic responses that may stay local or be propagated systemically through long-distance signalling (Motte *et al*., 2019; Ruffel *et al*., 2011; Ruffel *et al*., 2016). Growth inhibition under chemical stress does not necessarily result from irreversible cellular damage. In many cases, environmental constraints actively remodel meristem activity through coordinated developmental and transcriptional responses that transiently restrict cell proliferation (Motte *et al*., 2019). Whether biuret-induced root growth inhibition reflects passive toxicity or a regulated and reversible developmental response nevertheless remains unknown.

Here, we combined physiological, developmental, genetic, and transcriptomic approaches in *Arabidopsis thaliana* to investigate plant responses to biuret. We characterized the effects of biuret on root and shoot development, root meristem activity, and auxin and cytokinin signalling. Our forward genetic approach identified a biuret-resistant mutant, providing insight into the genetic basis of biuret sensitivity and demonstrating that biuret induces a reversible developmental response accompanied by major changes in plant growth and hormonal signalling.

## Results

### Biuret reversibly inhibits primary root growth and impairs shoot development

Biuret molecules present in urea formulation has negative effect on leaves of certain species (Achor and Albrigo, 2005). However, the effect of this molecule on root development has never been studied. In order to describe this effect we studied the primary root development of *A.thaliana* seedlings on media containing different concentrations of biuret. Plants were germinated and grown in presence of four different concentrations of biuret. The primary root length was measured over 12 days. Starting from 6 das, biuret induced a significant (*p* < 0.001) decrease in primary root growth (Figure 1 a,b). The observed effect was dose-dependent, since increasing the concentrations from lead to a reduction in PR length up to 98% at 12 das (Figure 1b). Conversely, concentrations lower than 0.5mM do not have significant impact on PR growth (Figure S1). Due to the presence of the biuret in ureic fertilizers, plants are usually exposed to this molecule in association with urea. When we studied the biuret effect on PR length in presence of 2.5 mM urea, we did not observe any significant difference, indicating that the presence of urea does not alter the effect of biuret (Figure S1). The root growth is dependent on the root apical cell division and elongation. To determine if the observed phenotype is due to a diminished cell division or the elongation, we used pWOX5::GFP quiescent center specific marker (Blilou *et al*., 2005) transgenic plants for assessing the number of cell composing the meristematic zone. The number of cortical cells in the meristem was assessed by counting on confocal microscopy images, in plants grown in presence of 1 mM biuret or control media w/o biuret. The results obtained show that biuret induce a 60% reduction in cortical cell number comparing to the control condition (Figure 1c,d). These results indicate that biuret inhibits root development mainly through a reduction in meristematic cell production, leading to a smaller meristematic zone. The effect of biuret on meristematic cell division activity was further investigated by analyzing the promoter activity of the cell cycle reporter gene *CYCB1;1* (Colón-Carmona *et al*., 1999). Transgenic reporter plants bringing the pCYCB1;1::GUS construct were cultivated on basal half-strength MS media and then transferred on media containing. The GUS staining indicates that high concentrations of biuret significantly decrease pCYCB1;1 promoter activity comparing to the control condition (Figure 1e). This result indicates that biuret inhibit the normal root development and this is associated to a decreased meristematic cell cycle activity in the root tip.

**Fig. 1.**
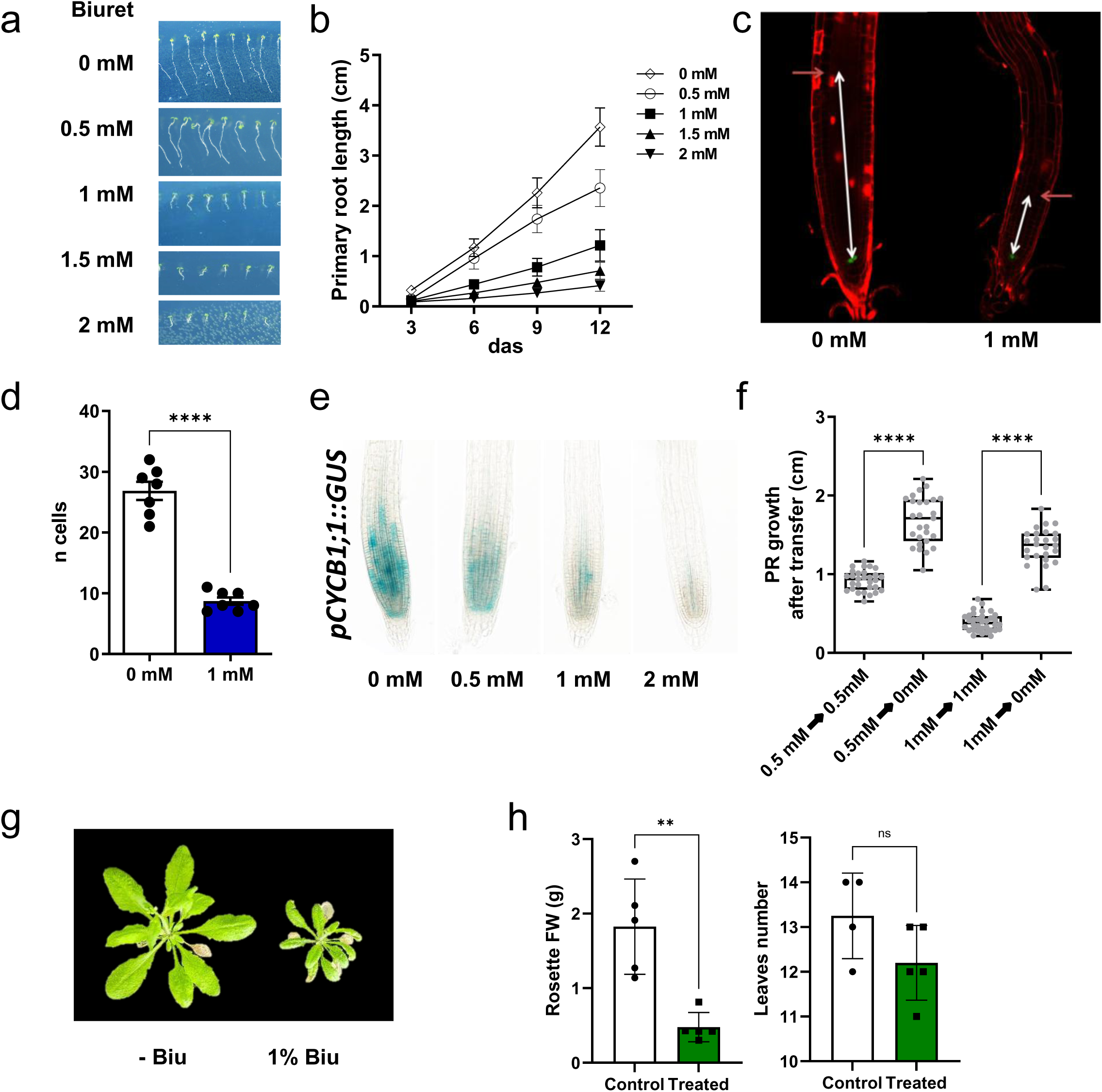
Biuret affects root and shoot growth. (a) Effect of biuret on primary root (PR) length 6 days after sowing. (b) Primary root length measured every 3 days in seedlings grown on media containing different biuret concentrations over a 12-day period. Data represent mean ± SD (n = 15–35). (c) Confocal images of pWOX5::GFP seedlings grown in the presence of 0 or 1 mM biuret and stained with propidium iodide (PI), showing differences in root meristem size. (d) Quantification of cortical cell number in the root meristem. The meristematic zone was defined as the region between the QC (identified by the WOX5:GFP signal) and the first elongated cortical cell. Asterisks indicate significant differences (\*\*\*\**p* < 0.0001; Student’s t-test). (e) Histochemical staining of pCYCB1;1::GUS seedlings grown for 6 days on media containing 0, 0.5, 1 or 2 mM biuret. (f) Length of the root segment produced during the 3 days following transfer from medium containing 0.5 or 1 mM biuret to biuret-free medium. Asterisks indicate significant differences (\*\*\*\**p* < 0.0001; one-way ANOVA followed by Tukey’s post hoc test). (g) Effect of repeated foliar spraying with a 1% (w/v) biuret solution on rosette growth. (h) Quantification of the effects of foliar biuret treatment on rosette fresh weight and leaf number. Three-week-old plants were sprayed daily with 1 mL of biuret solution or water (control; supplemented with 0.05% Silwet) for 14 days. Asterisks indicate significant differences (\*\**p* < 0.01; Student’s t-test), whereas ns indicates no significant difference.

Moreover, to verify if biuret lead to a complete and permanent exhaustion of the PR meristem activity, we made a transfer experiment where plantlets grown for 6 days in presence of 0.5 or 1mM biuret where transferred for 3 days on a new media without biuret. The results indicate that biuret does not irreversibly impair root meristem activity, as primary root growth rapidly resumed after transfer to biuret-free medium (Figure 1f).

The negative effects of biuret on leaves are largely documented (Mikkelsen, 1990). In order to characterize the impact of biuret on Arabidopsis shoot growth in our conditions, we made a phenotyping of the rosette in plants grown in pots and exposed to biuret foliar pulverization. We observed a clear inhibition of rosette growth in plants exposed to 1% biuret treatment (Figure 1g), characterized by a significant reduction in fresh weight and a slight, although not statistically significant, decrease in leaf number (Figure 1h). These observations suggest that biuret exerts a more pronounced effect on leaf expansion than on leaf emergence.

Taken together, these results, obtained using different experimental approaches on both roots and shoots, suggest that: (i) biuret strongly inhibits primary root (PR) growth in *Arabidopsis thaliana* in a dose-dependent manner by reducing meristematic cell division rather than cell elongation, an effect that is largely local and reversible; and (ii) biuret also negatively affects shoot growth by limiting leaf expansion.

### Biuret interferes with auxin and cytokinin signaling at the root apex

Auxin and cytokinin play central and antagonistic roles in the regulation of root meristem activity and root development (Overvoorde *et al*., 2010). As auxin distribution and response pattern play a central role in root development and growth, we investigated a possible role of this hormone in biuret inhibition of the PR growth. For that, we used DR5::GUS (Ulmasov *et al*., 1997) and DR5::GFP (Benková *et al*., 2003) reporter lines to analyze the effect of biuret on auxin signaling at the root tip. The DR5::GUS reporter revealed a marked alteration of auxin response patterns in the root apex of plants grown in presence of 2mM biuret. The GUS staining was detected as expected in columella cells, but in presence of biuret, the staining intensity was increased and extended to the stele cells (Figure 2a). Using the DR5::GFP reporter line we were able to quantify the promoter response to biuret, revealing a dose-dependent increase in DR5 reporter activity in response to biuret treatment (Figure 2b, Figure S2). The signal quantification showed that biuret induces a significant increase of auxin signal in the columella zone (+75-88% - ROI1) and the elongation zone (+50-150% - ROI2) in presence of 1 and 2mM biuret respectively (Figure 2c).

**Fig. 2.**
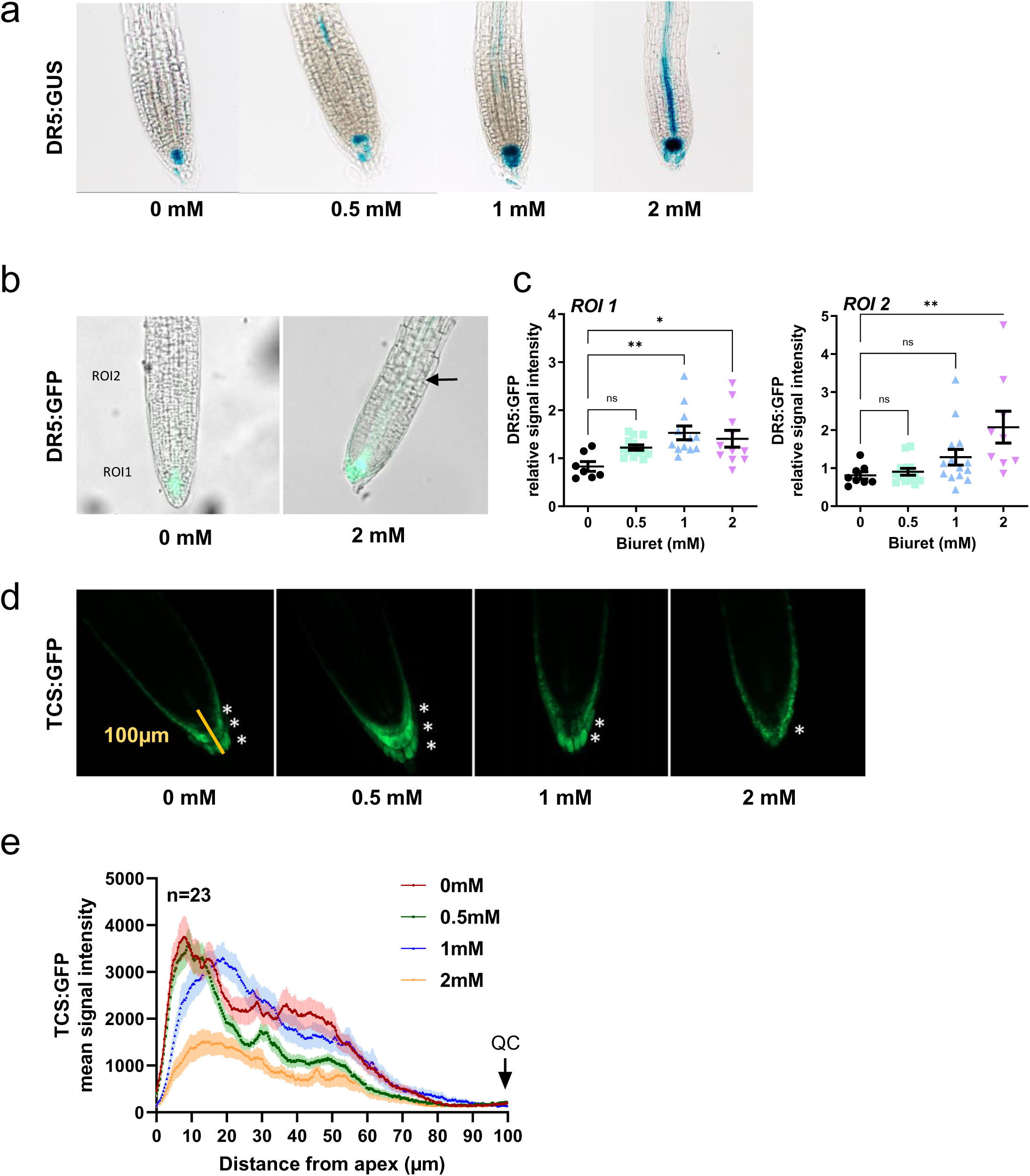
Biuret alters auxin and cytokinin signaling in the root. (a) Histochemical staining of the auxin reporter DR5::GUS and (b) fluorescence microscopy images of the DR5::GFP reporter line in seedlings grown for 6 days on media containing 0–2 mM biuret. (c) Quantification of DR5::GFP fluorescence intensity in two regions of interest (ROIs): the root division zone (ROI 1) and the elongation zone (ROI 2) (see Figure S3). (d) Fluorescence microscopy images of the cytokinin reporter TCS::GFP in seedlings grown for 6 days on media containing 0–2 mM biuret. White asterisks indicate cell layers displaying detectable TCS::GFP signal. (e) Quantification of TCS::GFP fluorescence intensity along a 100-µm transect extending from the root apex to the quiescent center (QC). Asterisks indicate significant differences relative to the 0 mM control condition (\**p* < 0.05, \*\**p* < 0.01, \*\*\**p* < 0.001; one-way ANOVA followed by Dunnett’s multiple-comparison test).

Cytokinins regulate not only the radial patterning and the cells proliferation but also contribute to the regulation of the root development in the longitudinal sense. Also, cytokinins control meristem size by directly regulating the auxin activity (Wybouw and De Rybel, 2019). To analyze the effect of biuret on cytokinins signaling, we studied the TCS::GFP reporter line (Zürcher *et al*., 2013) reporting cytokinin-responsive transcriptional activity. In absence of biuret, we observed the expected GFP signal in three columella cell layers, but the presence of biuret in the media (0.5 mM to 2mM) induces a perturbation in the cytokinin signal distribution. In fact, we observed a loss of fluorescence in columella cells and in particular, the ones laying just below the quiescent center, in presence of 1 mM and 2 mM biuret (Figure 2d). In the highest biuret concentration, the cytokinin signal was also lost in the outer columella cell layers. The GFP signal quantification confirmed this loss of fluoresce in different cell layers (Figure 2e), markedly evident in 2mM condition, consistent with a reduction in cytokinin-responsive activity rather than a loss of columella cell layers, whose number remained unchanged (data not shown).

Together, these observations, obtained on reporter lines in which the biuret phenotypes were consistent to the wild type (Figure S3), reveal that biuret profoundly perturbs the hormonal landscape of the root apex, with opposite effects on auxin- and cytokinin-responsive outputs. Interestingly, biuret treatment was also associated with callose deposition around the quiescent center region (Figure S4), suggesting that biuret-induced hormonal perturbations may be accompanied by modifications in symplastic connectivity at the root apex.

### Forward genetic screening identifies a biuret-resistant mutant

To identify genetic determinants controlling biuret sensitivity, we performed a forward genetic screen using approximately 10,000 Arabidopsis T-DNA insertion lines (Alonso *et al*., 2003) grown in media containing 1 mM and 2 mM biuret. Mutants displaying sustained primary root growth under inhibitory biuret concentrations were selected as putative biuret-insensitive lines. Because biuret-resistant phenotypes were readily identifiable, we visually selected 37 lines displaying longer primary roots than the wild type when grown in the presence of biuret and subsequently measured their primary root (PR) lengths. The wild-type (Col-0) genotype, used as an internal control, exhibited a shorter PR length (1.43 ± 0.03 cm) than the average PR length of the 37 selected mutant lines grown on 1 mM biuret (1.61 cm) (Figure S5a), thereby confirming the reliability of the visual screening procedure. The analysis allowed the identification of 8 lines with PR lengths significantly longer than the WT on 1mM biuret and two of them significantly longer on 2 mM biuret (Adj.*p*<0.05, Table S1). Among these eight, we selected three T-DNA insertion lines (# 23, 29, 36) with the strongest phenotype (Adj. *p*<0.0001) on 1mM biuret that we named *bir* (for <u>bi</u>uret resistant).

We further analyzed the root growth of *bir23*, *bir29* and *bir36* in presence of 0, 1 or 2 mM biuret (Figure S5b). *bir29* was the only mutant line showing a PR length significantly different in presence of 1 and 2 mM biuret. The effect on the PR root development was specific to biuret, since in control condition the PR was even shorter that the control genotype. Upon further validation, *bir23* showed a biuret insensitive phenotype only at high concentration (2 mM) and *bir36* showed no phenotype in presence of biuret but longer PR in the control condition. Considering these results, only *bir29* was retained for further analysis.

In control conditions, the *bir29* mutant was not different from the WT; but when grown in presence of biuret *bir29* mutant had significantly longer PR and increased LR number. In the presence of 1 mM and 2 mM biuret, the *bir29* mutant exhibited primary roots that were approximately 2-fold and more than 3.5-fold longer than those of the wild type, respectively (Figure 3a-b). Interestingly, the LR number was more than two-fold higher than the wild type in presence of biuret (1 mM and 2 mM)(Figure S5c). Moreover, we studied the effect of the mutation on the shoot development in control conditions and under foliar biuret treatment (biuret 1 %w/v). We observed that the mutant had different rosette biomass (FW) in both control and treated conditions (Figure 3c) but equal number of leaves (Figure 3d). In fact, comparing to the WT, *bir29* showed a significant reduction in rosette fresh weight in control (−37 %) and treatment (−3 %) conditions. Interestingly, the growth inhibition under treatment was 5 % less important in *bir29* than the WT. We also observed that, cultivated in media containing low concentrations of biuret (0.5 mM) that allow the rosette growth up to the floral transition, WT plants showed an inhibition of the inflorescence development (Figure 3e,f). In contrast, the *bir29* mutant developed the inflorescence even in presence of 0.5mM biuret. These results indicate that the genotype of *bir29* has strong effect both on root and shoot development in response to local biuret supply.

**Fig. 3.**
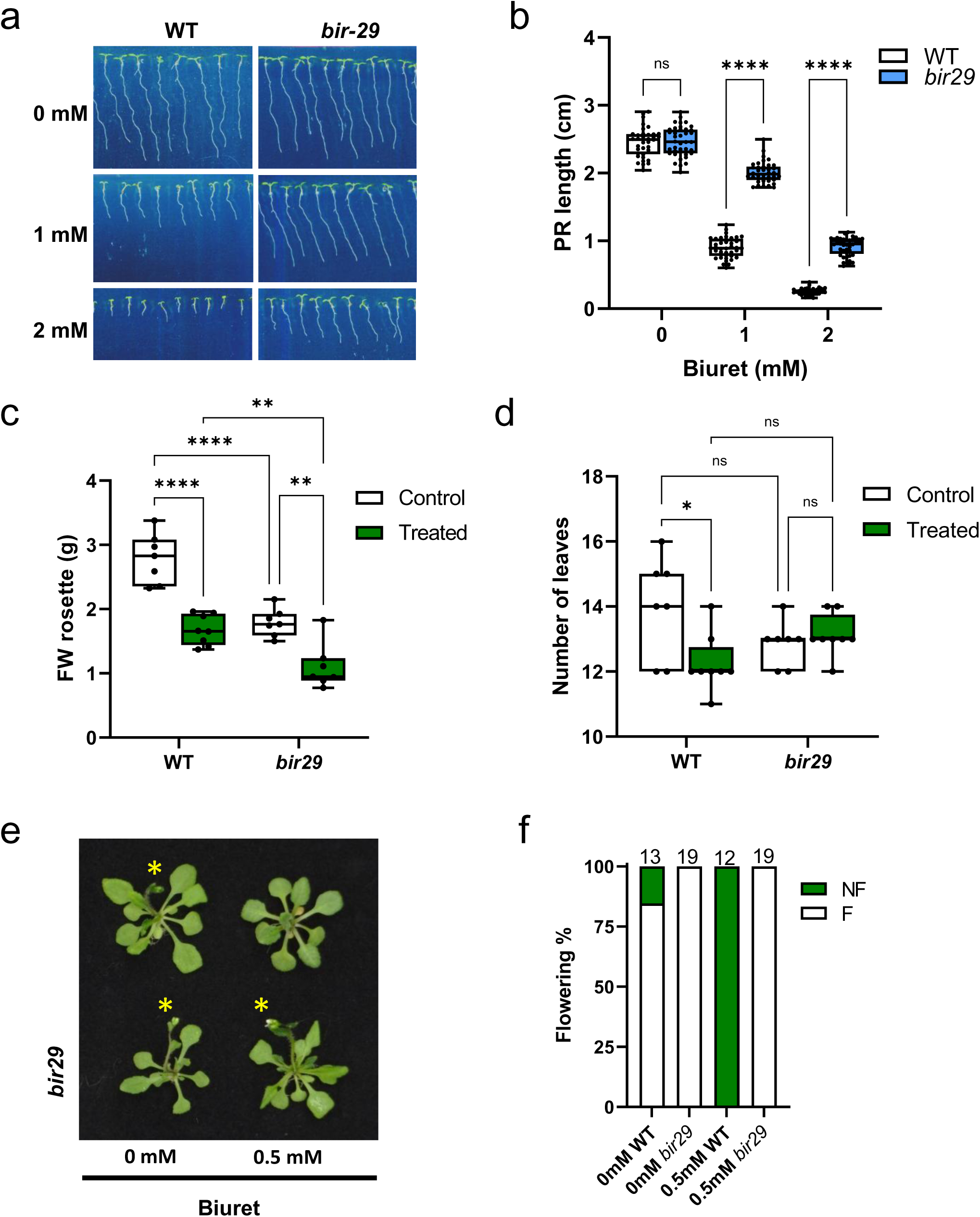
The *bir29* mutant is resistant to the inhibitory effects of biuret. (a) Representative root phenotypes of wild-type Col-0 (WT) and *bir29* seedlings grown under the same conditions. (b) Primary root length of WT and *bir29* seedlings grown for 6 days on media containing 0, 1 or 2 mM biuret. (c,d) Effect of a 14-day foliar treatment with a 1% (w/v) biuret solution on rosette fresh weight c) and leaf number d) in WT and *bir29* plants. (e) Representative images illustrating the effect of biuret supplied to roots on inflorescence development in WT and *bir29* plants. Yellow asterisks indicate inflorescences. (f) Percentage of flowering (F) and non-flowering (NF) plants after 4 weeks of growth *in vitro* on half-strength MS medium with or without 0.5 mM biuret. The total number of plants analyzed is indicated above each bar. Asterisks indicate significant differences (\**p* < 0.05, \*\**p* < 0.01, \*\*\*\**p* < 0.0001; two-way ANOVA followed by Tukey’s post hoc test).

### Altered auxin and cytokinin signalling underlies the biuret-resistant phenotype of *bir29*

To further investigate the relationship between the *bir29* phenotype in response to biuret and auxin/cytokinin signalling, we introduced the DR5::GFP and TCS::GFP reporters into the *bir29* genetic background, generating the DR5::GFP(*bir29*) and TCS::GFP(*bir29*) lines. These reporter lines were subsequently grown on media supplemented with different concentrations of biuret. Measurements of primary root (PR) length confirmed the resistant phenotype previously observed in the *bir29* mutant (Figure S3). Quantification of GFP fluorescence revealed an increase in the DR5 response (+36%) in the *bir29* background upon treatment with 2 mM biuret. This increase was specifically observed in the columella region (ROI1) (Figure 4b), whereas the signal remained unchanged in the stele (ROI2) under all tested conditions (Figure 4a–b). Consistently, analysis of the TCS signal showed that fluorescence remained detectable across the three columella layers (Figure 4c), even under high biuret conditions (2 mM). Quantification revealed an increase (+40%) in the *bir29* background (Figure 4d).

**Fig. 4.**
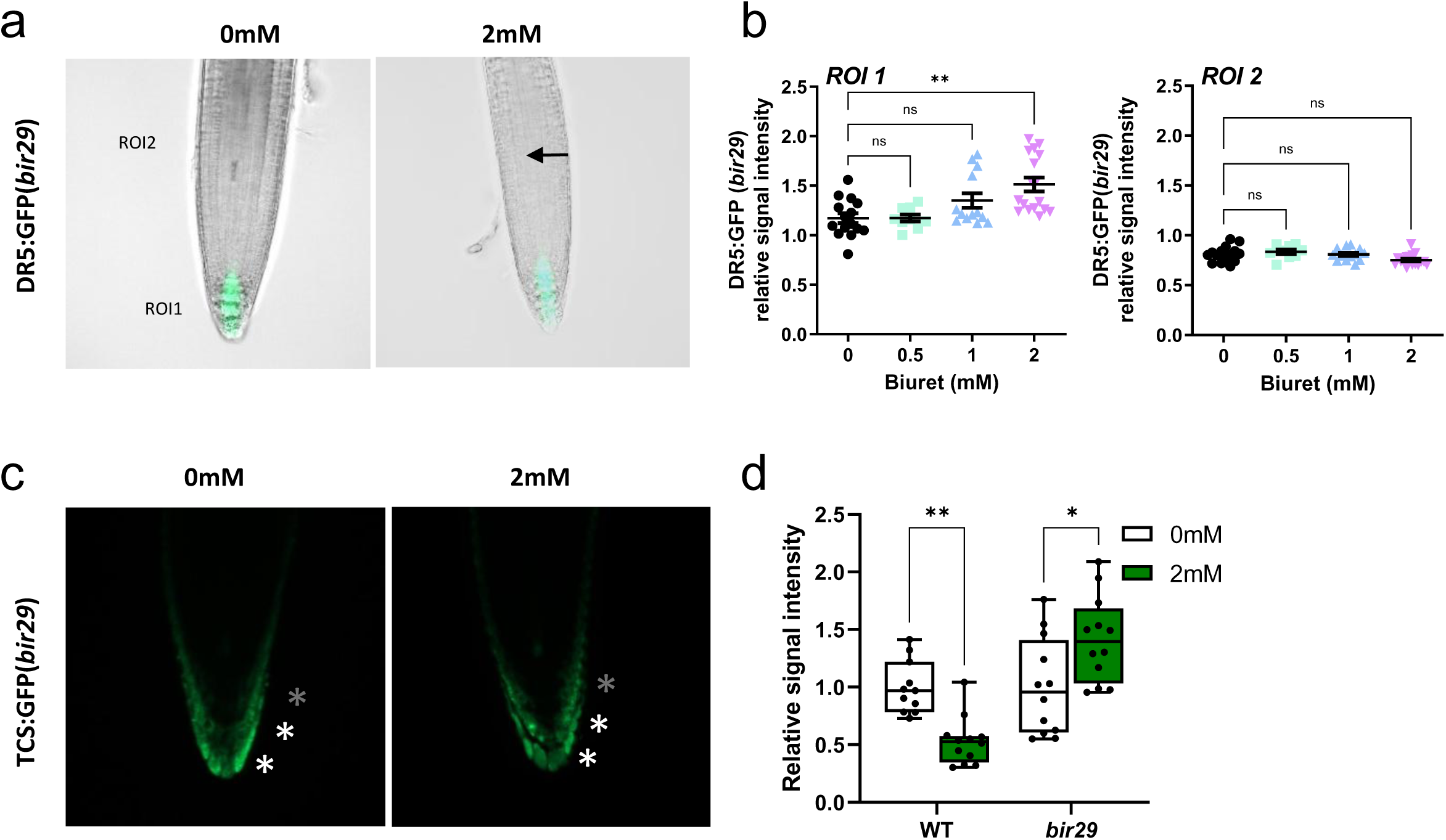
Biuret-induced changes in auxin and cytokinin signaling are altered in the *bir29* mutant. (a) Fluorescence microscopy images of the auxin reporter DR5::GFP in the *bir29* genetic background in seedlings grown for 6 days on media containing 0–2 mM biuret (see Figure S3 for intermediate concentrations).(b) Quantification of DR5::GFP fluorescence intensity in two regions of interest (ROIs): the root division zone (ROI 1) and the elongation zone (ROI 2). Values are expressed relative to the mean fluorescence intensity of the 0 mM condition. (c) Fluorescence microscopy images of the cytokinin reporter TCS::GFP in the *bir29* genetic background in seedlings grown for 6 days on media containing 0–2 mM biuret. White asterisks indicate cell layers displaying detectable TCS::GFP signal. (d) Quantification of TCS::GFP fluorescence intensity along a 60-µm transect extending from the root apex to the first columella layer in WT and *bir29* roots. Values are expressed relative to the mean fluorescence intensity of the 0 mM condition. Asterisks indicate significant differences between WT and bir29 under the same treatment condition (\**p* < 0.05, \*\**p* < 0.01, \*\*\**p* < 0.001; two-way ANOVA followed by Tukey’s post hoc test).

Taken together, these response patterns differ from those observed in the wild-type background (Figure 2; Figure 4d), suggesting that in the *bir29* mutant, auxin and cytokinin responses upon biuret treatment are altered. This alteration could represent either a cause or a consequence of the bir29 phenotype associated with the underlying genetic mutation.

### Biuret triggers a systemic effect on primary root growth that is impaired in *bir29*

It has been described that chemicals can be sensed locally and induce a systemic root development remodeling through long distance signaling pathways (Ruffel *et al*., 2016). We tested the local and systemic effect of biuret on root growth. Our split-root experiments (Figure 5a, b) revealed a strong inhibition of root growth in the locally treated compartment (Sp Biu2 vs Sp Biu0) and a significant inhibition was also detected in the untreated compartment (Sp Biu0 vs C Biu0). This suggests that biuret response is predominantly local but also involves a systemic component. In contrast, to the strong primary root phenotype, lateral root development was only marginally affected under our experimental conditions (Figure S6a,b,c). Interestingly, the systemic response to biuret in *bir29* mutant was modified since we found no significant difference for the PR length between the control and the split conditions (Figure 5a). In contrast, the systemic effect on LR development was not altered in *bir29* (Figure S6a,b,c). These results underlie the signaling behavior and the genetic control of the root biuret response.

**Fig. 5.**
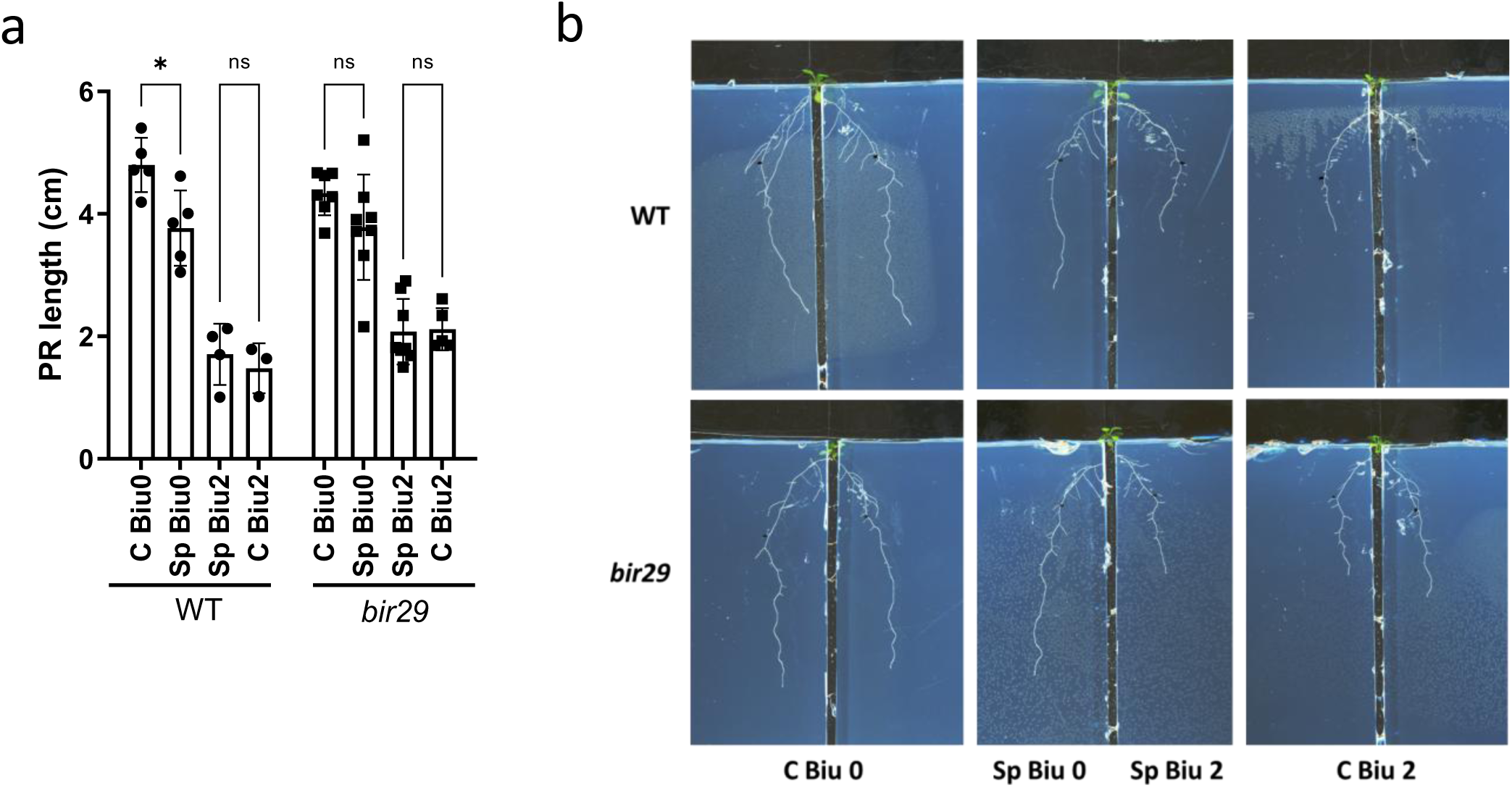
Local and systemic effects of biuret on root development in the *bir29* mutant. Quantification of primary root length (a) in wild type and *bir29* plants grown in a split-root system. (b) Photos of the roots of plants cultivated in the split root system. Seedlings were initially grown on basal medium and subsequently transferred to the split-root system for 6 days. C Biu0 and Sp Biu0 correspond to biuret-free conditions, whereas Sp Biu2 and C Biu2 contain 2 mM biuret. In control plants (C), both root halves were exposed to the same medium, whereas in split plants (Sp), the two root halves were exposed to different media. Asterisks indicate significant differences (*p* < 0.05; one-way ANOVA followed by Tukey’s post hoc test).

### The biuret-resistant phenotype of *bir29* is not caused by reduced uptake of the molecule

In rice, biuret has been reported to be absorbed through the root system (Ochiai *et al*., 2020). To determine whether Arabidopsis plants can absorb biuret through their roots and whether the biuret-resistant phenotype of *bir29* results from a reduced uptake rate or a lower long-term accumulation of biuret (reflecting the balance between influx and efflux), we investigated biuret uptake and its accumulation in roots and shoots using ^15^N-labeled biuret. We observed that WT and *bir29* had similar influx rate (14.06 and 12.02 µmol.gDW^-1^.h^-1^ respectively) when 1 mM biuret was supplied (Figure 6a). Moreover, in three days plants whose roots were exposed to 0.5 mM biuret accumulated ^15^N-biuret (or a derived ^15^N-containing molecule) up to 22-24 µmol.gDW^-1^ in the roots and up to 18-19 µmol.gDW^-1^ in the shoots (Figure 6b). For both, rapid influx (5 minutes) and long-term (three days) accumulation, we did not observe any significant difference between the WT and the mutant, indicating that the resistant phenotype of *bir29* is independent of biuret uptake and internal translocation.

**Fig. 6.**
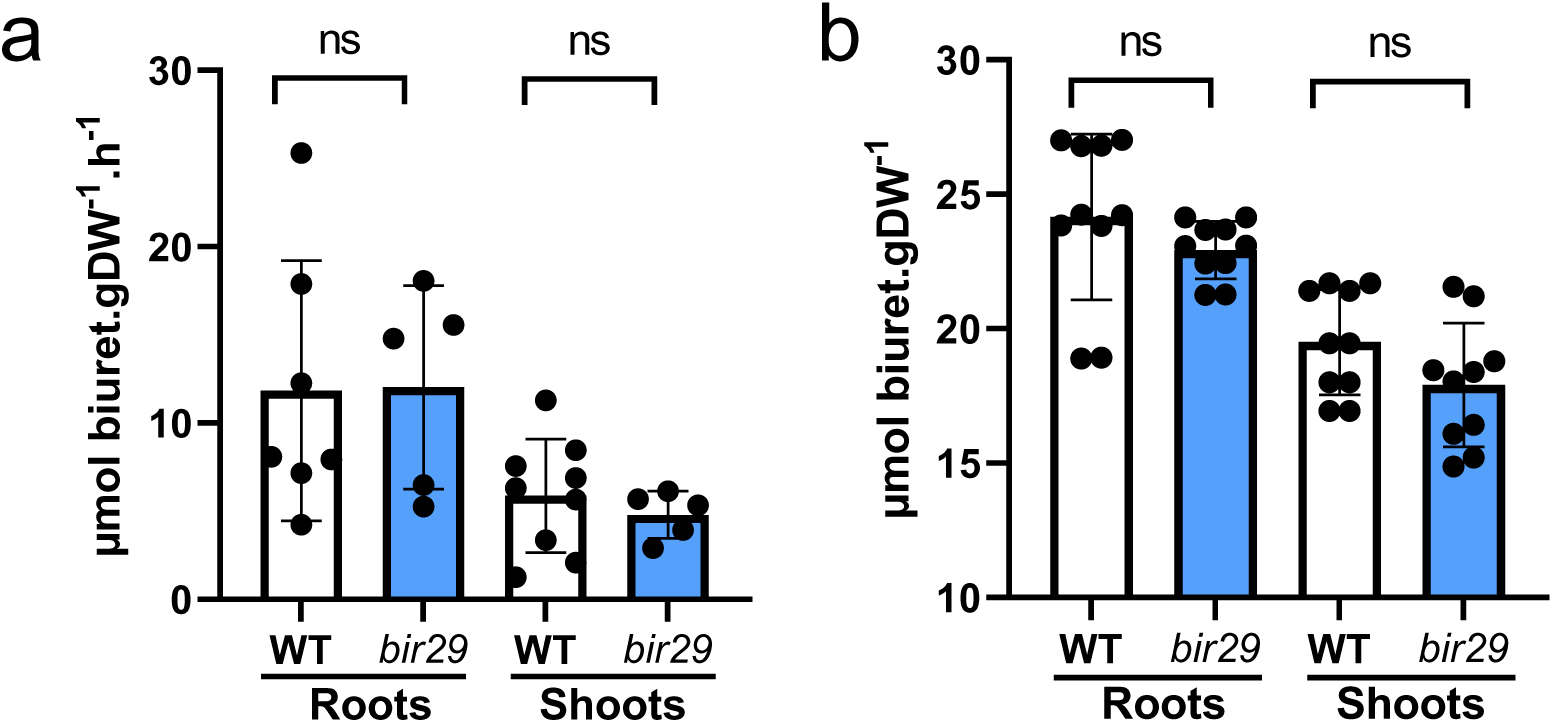
Biuret accumulation and influx do not differ in WT and *bir29* plants. (a) Biuret influx rates in roots and shoots of twelve-day-old WT and *bir29* seedlings grown *in vitro*. Roots were immersed in 1 mM ^15^N-labelled biuret for 5 min before isotope quantification. (b) Biuret accumulation in roots and shoots of four-week-old WT and *bir29* plants grown in a hydroponic system. Plants were exposed to a 0.5 mM solution of ^15^N-labelled biuret added to the nutrient medium for 3 days. Bars indicates means ±SD. ‘ns’ indicates no significant difference between WT and *bir29* (unpaired Student’s t-test).

### The genome wide transcriptional response of *bir29* mutant to biuret reveals potential mechanisms

To obtain further insight into the root response to biuret supply and its relationship with the *bir29* mutant phenotype, we performed a transcriptomic analysis on three independent experiments. For that, roots tissues of WT and *bir29* mutant cultivated for 6 days in control or biuret containing (1mM) media were sampled and analyzed. As the objective of this analysis was exploratory, the parameters were intentionally set with low stringency. Data were modeled as previously described (Medici *et al*., 2015)(see methods for details). Analysis of variance (ANOVA) retrieved 3233 and 2008 genes whose expression was significantly regulated (*p* <0.05) by the biuret treatment and the genotype respectively (Table S2-S3), what highlights the strong effect of both factors independently. In contrast, only a small set of 858 genes was significantly regulated by the genotype*treatment interaction (Tukey adjusted *P* < 0.05; Table S1, S4). We performed hierarchical clustering of both genes and samples of these 858 genes and we first observed the similarity between the gene expression profiles in WT control samples and both control and biuret-treated *bir29* samples (Figure 7a), the biuret-treated WT samples laying in a distinct branch on the left side of the dendrogram. This distribution closely resemble the phenotypic patterns observed for WT and *bir29* mutants (Figure 3b). The H-Clustering of the 858 differentially expressed genes (DEGs), identifies four main clusters (C1 to C4). C1 and C3 contain genes that are strongly upregulated (C3) and downregulated (C1) in the WT in response to biuret, but showing weak variation in *bir29*. C4 contains genes strongly downregulated in *bir29* by biuret that were not in WT; C2 corresponds to genes that have opposite regulation in WT (down) and *bir29* (up) in response to biuret treatment. The 858 DEGs consisted of nearly equal numbers of upregulated (log2FC > 0) and downregulated (log2FC < 0) genes in both genotypes, with 425/433 in WT and 427/431 in the *bir29* mutant (Figure 7b). To identify the genes potentially involved in the biuret-resistant phenotype, we studied the list of genes having highly different regulation between the two genotype. From the intersection analysis (Figure 7b, Table S5), groups 1 and 2 correspond to genes that have completely opposite regulation between the two genotypes and account for 65% of the total DEGs. In contrast, intersection groups 3 and 4 correspond to genes showing similar regulation (although likely with different magnitudes) in both genotype and account for only 35%. These results indicate a strong differential response between WT and the *bir29* mutant upon biuret treatment, which likely underlies the *bir29*-resistant phenotype. We conducted a GeneOntology analysis of the intersection groups and found significant overrepresentation of terms for 3 of them (Table 1, Table S6). The gene set of the 277 genes down-regulated in the WT and up-regulated in the mutant (Intersect group 1) was enriched in terms “microtubule-based movement” (GO:0007018); “meiotic cell cycle” (GO:0051321) and “sister chromatid segregation” (GO:0000819). Across these enriched GO categories (5- up to 13-fold enrichment, FDR<0.05), the repressed genes in the WT predominantly encoded kinesin (*ATK1*, *HIK*) - and cell cycle-associated proteins (Table 1 and Table S1), supporting the hypothesis that biuret exposure could suppresses microtubule-dependent cell division and root meristem activity in wild-type plants. In the corresponding intersection, the list of 275 genes upregulated in WT and downregulated in the mutant (intersection group 2) was enriched in the terms “response to oxygen-containing compound” (GO:1901700) and “water transport” (GO:0006833). Analysis of these genes revealed concomitant regulation of aquaporins (*PIP2;1, PIP2;2, PIP1;2, PIP1;5*, and *TIP2;2*), together with genes involved in reactive oxygen species (ROS) signaling and abiotic and biotic stress adaptation, including *RBOHF*, *CAMTA3*, *NDR1*, *IOS1*, *ERD15*, *LEA30*, *RAP2-4*, *SWEET17*, and *MYB88*. This suggests an integrated regulation of hydraulic conductance, redox signaling, and osmotic stress responses that is absent in the mutant. The list of 150 genes upregulated in both genotypes (intersection group 4) was enriched in the term “ribosomal large subunit biogenesis” (GO:0042273), highlighting a core translational regulation signature associated with increased expression of ribosomal proteins and assembly factors (*RPL7A*, *RPL7B*, *SMO4*, *EL33X*, and *EL8Y*). This suggests an activation of ribosome assembly and protein synthesis capacity in both genetic backgrounds in response to biuret treatment. Finally, no term was significantly enriched in the list of 156 genes downregulated in both genotypes (intersection group 3).

**Fig. 7.**
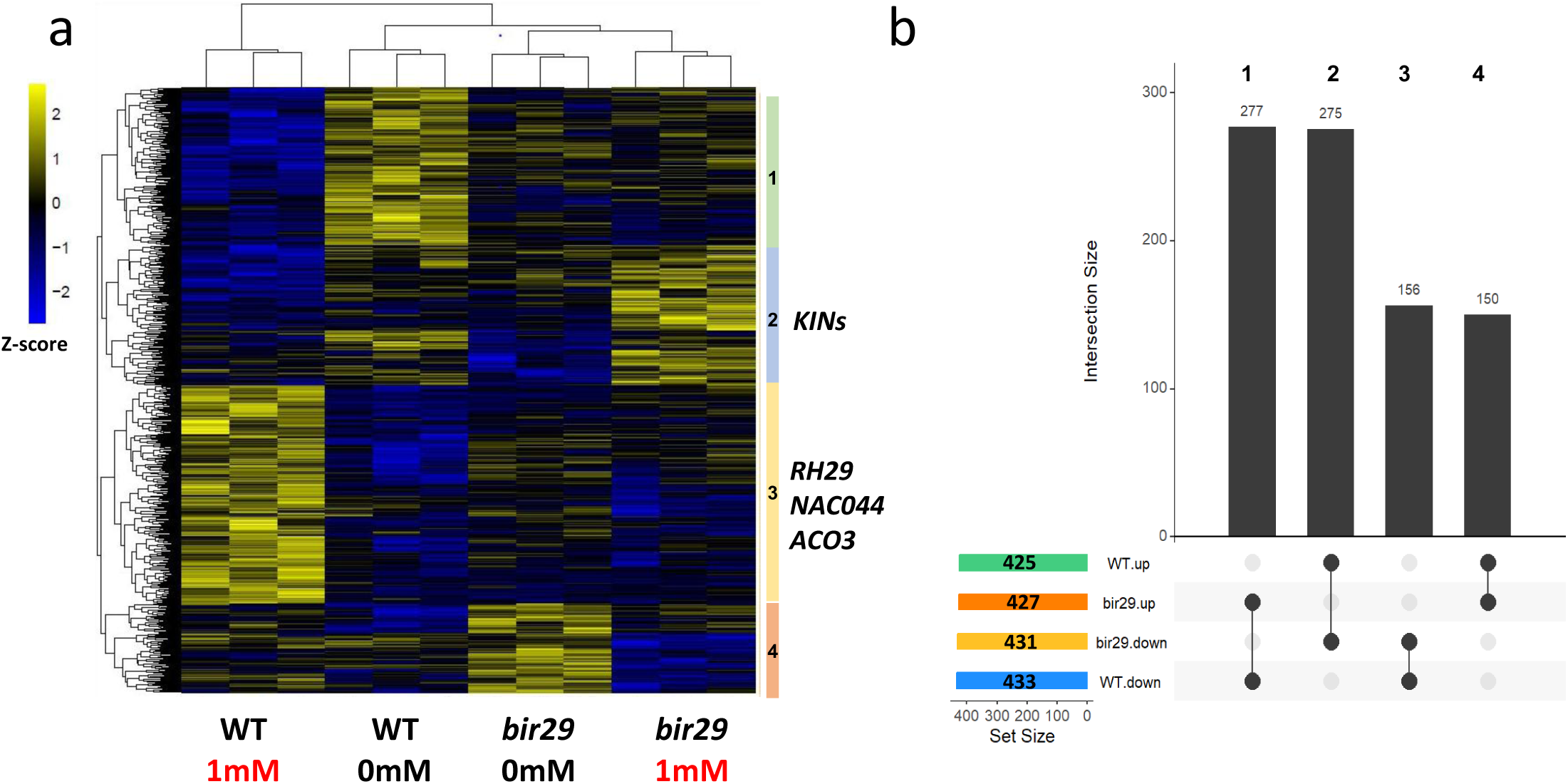
The genome-wide transcriptional response to biuret is strongly altered in the *bir29* mutant. (a) Heat map showing the centered and scaled expression values of the 858 genes differentially expressed in WT (Col-0) and *bir29* roots under biuret treatment (*adj. p* < 0.05). Hierarchical clustering of genes and samples is shown on the left and top sides of the heat map, respectively. Gene clusters displaying distinct expression profiles are indicated by colored labels and cluster numbers. (b) UpSet plot showing the intersection sizes among four groups of genes displaying shared or contrasting transcriptional responses to biuret in WT and *bir29* plants. Genes with log2FC > 0 were considered up-regulated, whereas genes with log2FC < 0 were considered down-regulated. Intersection identifiers corresponding to the data reported in Table 1 are indicated above each bar.

These results suggest that biuret is absorbed and perceived at the root level, inducing a specific response likely associated with the production of reactive oxygen species and triggering a signaling cascade involving components of the pathogen-triggered immunity for instance, since the genes associated with these GO terms are upregulated only in WT. Moreover, the fact that genes involved in cell cycle processes are downregulated in WT but not in the *bir29* mutant is consistent with the reduced activity of the CYCB1 marker observed in response to the treatment (Figure 1e) and with the stunted growth phenotype.

To go further we look in the list of the 858 DEGs for genes that had high expression regulation - |log2FC|>1 (Table S7). For the genes strongly regulated in WT and loosing this regulation in *bir29* in response to biuret (laying in Cluster 3 Figure 7a), we found some interesting (Figure S8a). *ACO3* (*AT1G12010*), for example, showed a more than two-fold induction in WT root in presence of biuret, but this regulation was loss in *bir29* (Cluster 3). *ACO2*, *ACO3*, and *ACO4* are responsible for ethylene biosynthesis and ethylene-regulated root and RAM shortening in cytokinin-treated *Arabidopsis* (Yamoune *et al*., 2024). The *LURP1-like DUF567* gene (*AT1G33840*), known to be expressed in response to oomycete infections (Knoth and Eulgem, 2008), showed a three-fold induction in the WT, that was reduced to 0.08-fold change in the mutant. The same pattern was found for the RNA helicase *RH29* (*AT1G77030*) (Chen *et al*., 2020) whose upregulation was two-fold higher in the WT than in *bir29* and the NAC-type transcription factor *NAC044* (*AT3G01600*), required for DNA damage-induced G2 arrest under genotoxic stress conditions (Takahashi *et al*., 2019), whose three-fold expression induction was lost in *bir29*.

Moreover, two genes induced by biuret in the wild type were constitutively expressed at high levels in *bir29*. The first was *BGLU26/PEN2* (*AT2G44490*), a defense-related β-glucosidase involved in indole glucosinolate-mediated immunity (Lipka *et al*., 2005). The second was the non-coding RNA *AT4G04223*, previously reported to be upregulated in the roots of the *xal2-2* mutant, which exhibits a pronounced short-root phenotype (Castañón-Suárez *et al*., 2024) (Figure S8b). Conversely, three genes were specifically upregulated in *bir29* (Figure S8c). Among them, *AT2G21640* has been described as a marker of oxidative stress responses and is induced by sucrose and the herbicide atrazine (Ramel *et al*., 2007). The two others, *AT2G03230* and *AT3G54530*, encode a GSK domain-containing protein and a hypothetical protein, respectively, and remain poorly characterized. Overall, these results suggest that the *bir29* mutation reshapes the transcriptional response to biuret, leading to a partial uncoupling of stress, immune, and developmental signaling pathways.

### Genetic characterization of *bir29* reveals two T-DNA insertions associated with biuret resistance

The *bir29* mutant was identified through a forward genetic screen, using the Salk T-DNA insertion collection (Alonso *et al*., 2003). To characterize the genetic basis of the mutant phenotype we crossed *bir29* and the wild type. The resulting F₂ population, displayed an intermediate and highly variable distribution of relative primary root length compared with the parental lines (Figure 8a).The frequency distribution of the F_2_ (Figure 8b) shows a continuous unimodal and non-normal distribution (Shapiro-Wilk *p-*value = 0.0013), with a slight positive skew. The F_2_ population does not show values corresponding to the mean value of the *bir29* parental distribution (Relative PR length = 3.7), suggesting that the observed F_2_ distribution cannot be explained by a single gene. These results suggests an additive genetic effect of the *bir29* allele and the involvement of multiple loci rather than a single Mendelian locus that would generate a bimodal or multimodal distribution. To identify the T-DNA insertions present in *bir29*, whole-genome resequencing of *bir29* genomic DNA was performed. Sequencing reads were aligned to the *Arabidopsis thaliana* reference genome (Col-0 accession) and compared with reads obtained from a wild-type control. Genomic analysis conducted on the Arabidopsis TAIR10 assembly revealed that *bir29* mutant carries two T-DNA insertions. One T-DNA is inserted on chromosome 1 (position 18055357) within the coding sequence of *AT1G48820*, which encode a terpenoid cyclase protein - *TPS* (Chen *et al*., 2003). A second T-DNA insertion was identified on chromosome 3 (position 20560366) in the intergenic region between *AT3G55450*, encoding a PBS1-like protein serine/threonine kinase – *PBL1* - (Ranf *et al*., 2014; Tang *et al*., 2017) and *AT3G55460*, encoding an SC35-like splicing factor – *SCL30* - (Barta *et al*., 2010). This second insertion was precisely located 135bp and 274bp downstream the 3’UTR end of *PBL1* transcripts (Figure 8c) and 430bp upstream the 5’UTR beginning of *SCL30* transcript. Additionally this insertion was associated to an 11bp deletion (Figure 8c and S7b). Subsequently, PCR-based genotyping and fragment re-sequencing confirmed the two insertions and the deletion in these genomic loci (Figure S7a).

**Fig. 8.**
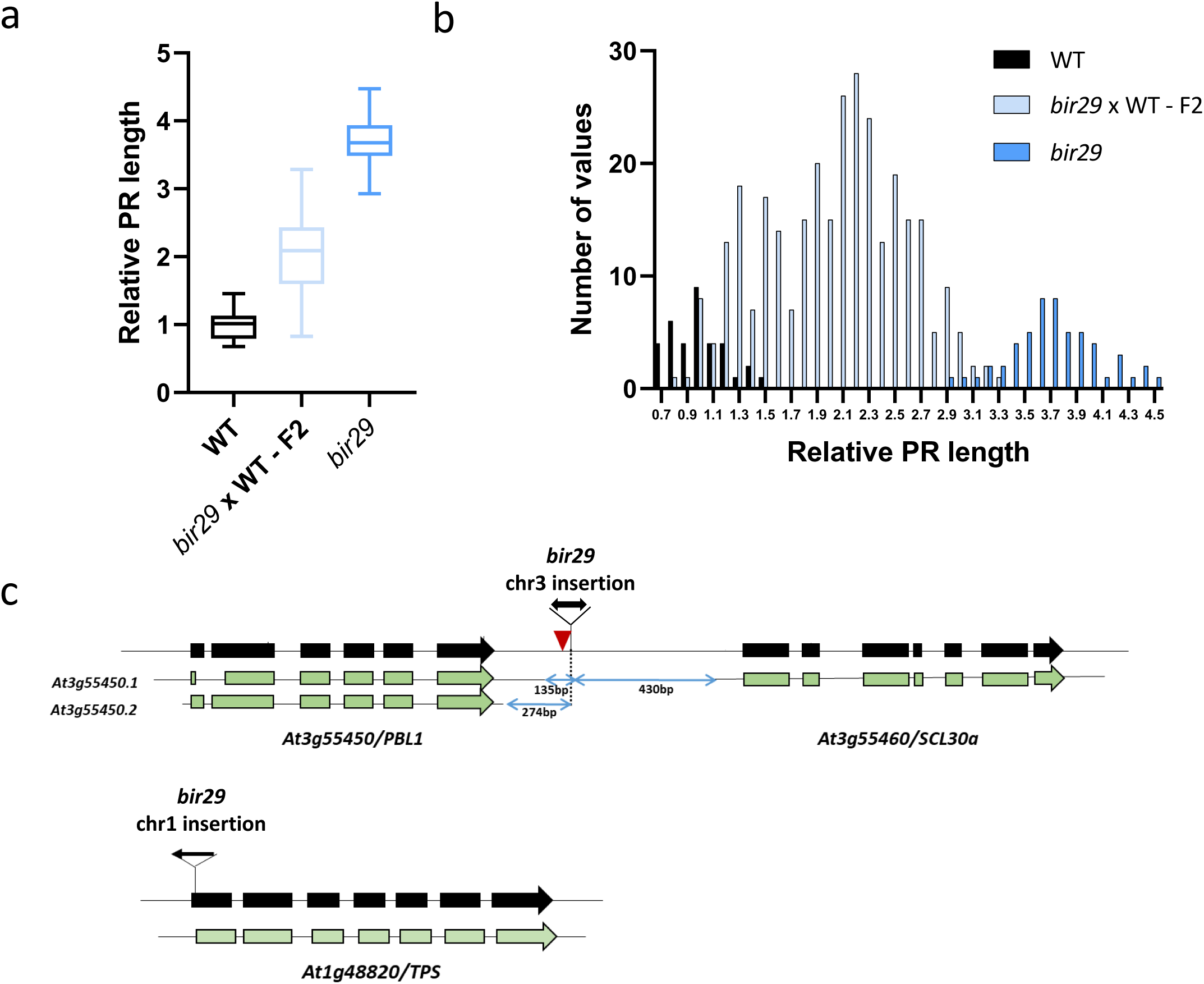
Multiple genomic loci involved in *bir29* phenotype. (a) Relative primary root length of wild-type (WT), the segregating *bir29* x WT F_2_ population and *bir29* seedlings grown on 1 mM biuret for 6 days. Primary root length was normalised to the WT mean. Box plots show the min to max distribution of individual measurements. (b) Frequency distribution of the relative primary root length values shown in (a) (bin width = 0.15). The colour code is the same as in (a). (c) Schematic representation of the T-DNA insertion sites (black arrows) identified in the *bir29* mutant. Insertions are located in the At3g55450–At3g55460 intergenic region and in At1g48820. Arrow orientation indicates the position of the T-DNA left border (LB). Alternative transcript isoforms of *PBL1* are shown. Blue arrows indicate the distance (bp) between each insertion site and the nearest UTR.

## Discussion

Biuret has long been regarded primarily as a phytotoxic contaminant of urea-based fertilizers (Achor and Albrigo, 2005; Mikkelsen, 1990), yet two of our observations are difficult to reconcile with a model based solely on toxic damage. First, biuret-induced inhibition of primary root growth is reversible, with growth resuming after transfer to biuret-free medium (Figure 1f). Second, the biuret-resistant mutant *bir29* maintains root and inflorescence development under inhibitory concentrations without any detectable change in biuret influx (Figure 3; Figure 5). Neither feature is readily explained by a model based solely on cumulative toxic damage. Together, these observations suggest that growth arrest involves active regulatory processes downstream of biuret exposure rather than resulting exclusively from cumulative toxicity, as would be expected for classical phytotoxins such as paraquat, whose redox cycling generates superoxide radicals that drive lipid peroxidation and progressive, irreversible damage to cellular membranes (Brunharo and Hanson, 2017). We propose that this reframing, from phytotoxic contaminant to a compound integrated into endogenous signalling networks, provides the interpretative framework through which the remaining results are best read.

At the cellular level, this reversibility maps onto the root meristem. Biuret reduced meristem size and cell-cycle reporter activity (Figure 1c-e), pointing to a controlled decrease in cell production rather than to an elongation defect, and proliferative competence was regained once the compound was removed (Figure 1f). The meristem is therefore more plausibly held in an actively regulated, growth-restricted state than driven to exhaustion. Such reversible regulation of meristem activity is a recurrent feature of plant responses to environmental constraints, salinity and nutrient limitation (Geng *et al*., 2013; Gutiérrez-Alanís *et al*., 2018; Sánchez-Calderón *et al*., 2005; West *et al*., 2004) in which growth is transiently restrained until favourable conditions return (Motte *et al*., 2019). Our findings place biuret within this broader class of perturbations that reshape developmental programmes through regulatory mechanisms and not solely through cytotoxicity.

At the root tip, this remodelling is read out as a coordinated shift in hormonal signalling: biuret enhanced auxin-responsive (*DR5*) output while reducing cytokinin-responsive (TCS) output (Figure 2). Because the auxin to cytokinin balance sets the boundary between proliferation and differentiation and thereby defines meristem size (Benková *et al*., 2003; Wybouw and De Rybel, 2019), this combined shift provides a plausible interpretative framework for the observed contraction of the proliferative domain. We emphasise, however, that DR5 and TCS report transcriptional outputs rather than hormone concentrations, and our data do not establish a causal link between these changes and growth inhibition; they are better viewed as hallmarks of a broader reorganisation of meristem function. In this context, the callose deposition detected around the quiescent centre (Figure S4) is of particular interest. Because callose can modify symplastic connectivity and the movement of mobile developmental regulators (Vatén *et al*., 2011), the formation of callose-rich domains around the stem-cell niche raises the possibility, to be tested functionally, that biuret reshapes the spatial organisation of signalling within the meristem. Whether this is a primary target of biuret or a downstream consequence of meristem reorganisation remains to be determined.

Biuret-induced responses are not confined to directly exposed tissues. In split-root experiments, growth inhibition was strong in compartments receiving biuret, but untreated roots also showed reduced growth relative to their controls (Figure 5), revealing a systemic component accompanying a predominantly local effect. Notably, this systemic component is lost in *bir29*, in which primary root growth no longer differs between control and split conditions (Figure 5), indicating that the mutation affects not only local sensitivity but also the long-distance dimension of the response. The nature of this systemic effect cannot, however, be resolved from the presented data. It may reflect a long-distance signal generated upon local biuret perception, but it may equally reflect biuret itself, or a nitrogen-rich derivative, taken up on the treated side and redistributed to the untreated roots. The marked accumulation of biuret-derived ¹⁵N in shoots (Figure 6b) is consistent with this second possibility, although redistribution to the untreated roots would additionally require phloem mobility, which the shoot data do not establish. A reduced supply of shoot-derived resources to all roots cannot be excluded either. Discriminating a generated signal from translocation of the compound would require quantifying ¹⁵N in the untreated compartment; we therefore limit our interpretation to the demonstrated existence of a systemic component, which places the biuret response within whole-plant rather than strictly local (Fu and Dong, 2013; Gilroy *et al*., 2014; Ruffel *et al*., 2011).

The hormonal set-point shift and the transcriptional repression of the cell-cycle machinery can be read as two outputs of the same meristem contraction. The isolation of *bir29* gives this interpretation a genetic basis. Because short-term biuret influx does not differ between wild type and mutant (Figure 6), resistance cannot be attributed to reduced entry; it acts downstream of uptake and separates biuret accumulation from biuret sensitivity. Strikingly, the resistant background bypasses the wild-type response to biuret. In our transcriptomic comparison, *bir29* neither represses the cell-cycle and microtubule machinery that biuret down-regulates in wild-type roots (intersection group 1; kinesins *ATK1* and *HIK*, and cell-cycle genes) nor deploys the redox, osmotic and stress-associated programme that biuret induces in the wild type (intersection group 2; *RBOHF*, *CAMTA3*, *NDR1* and *IOS1*, together with aquaporins and other water-transport genes), while the overall transcriptional state of biuret-treated *bir29* remains close to that of untreated wild type (Figure 7; Table 1). Because *bir29* does not arrest growth, the absence of these regulations could equally reflect a consequence of sustained proliferation rather than its cause. Several of the genes induced by biuret in the wild type but not regulated in *bir29* reinforce this reading: *NAC044*, required for DNA-damage-induced G2 arrest under genotoxic stress; *ACO3*, an ethylene-biosynthesis gene associated with cytokinin-dependent shortening of the root apical meristem; and the oomycete-responsive *LURP1*-like gene (AT1G33840), in line with the immunity-like component of the response (Figure S8a). The wild-type programme is therefore consistent with an active brake on proliferation and the interpretation of biuret-induced inhibition as a self-imposed restraint rather than chemical injury remains a hypothesis to be tested functionally: whether the stress- and defence-like component reflects bona fide immune engagement is still to be established.

What kind of resistance is this? From a genetic point of view, the continuous and non-normal distribution of phenotypic values in the F_2_ backcross population, the marked difference between the parental lines and the identification of the two T-DNA insertions (Fig 8a,b) are consistent with the idea that the biuret resistance is driven by multiple loci or a major-effect locus whose phenotypic expression is modified by additional genetic factors (at least one other insertion mutation). The fine characterization of these multiple loci is of high interest and requires extensive investigation, which is beyond the scope of the present work and will be addressed in a subsequent study. From a functional point of view, the reduced uptake can be excluded, since both the short-term influx and the longer-term accumulation of biuret are unchanged in *bir29* (Figure 6a,b), which also argues against active efflux. Three possibilities nonetheless remain open: i) total ¹⁵N does not distinguish intact biuret from nitrogen-rich derivatives, so intracellular detoxification cannot be excluded, ii) compartmentation of biuret into the vacuole, apoplast, or cell wall cannot be excluded and may reduce its effective cytosolic concentration, iii) because the molecular target of biuret is unknown, target-site insensitivity cannot be excluded either. What the data do establish is that resistance is uncoupled from biuret influx, since *bir29* sustains root and inflorescence growth while retaining as much biuret as the wild type. However, these three explanations are all local, and none of them easily explains why *bir29* also loses the entire wild-type transcriptional response and the systemic effect (Figure 5a). We therefore favour the hypothesis that *bir29* acts at the level of biuret perception and signalling, rather than in the metabolism or sequestration of the compound itself, via a complex multigenic mechanism.

A central question rise from this study. Land plants appear to lack a dedicated, efficient route to catabolise biuret, a compound that soil microbes consume as a nitrogen source (Ochiai *et al*., 2020; Robinson *et al*., 2018); one might therefore expect indifference or simple intoxication rather than a structured, signalling-dependent and genetically tractable response. Our data do not support biuret being read primarily as a nitrogen source. The few nitrogen-related genes differentially modulated between wild-type and mutant roots were the RING-type E3 ubiquitin ligase *NLA* (Nitrogen Limitation Adaptation), involved in plant adaptation to nutrient stress (Table 1), and the nitrilases *NIT1* and *NIT2*, which convert indole-3-acetonitrile to indole-3-acetic acid (IAA) (Table S3). Given the described role of NLA in integrating different environmental constraints (Kant *et al*., 2011; Park *et al*., 2014; Val-Torregrosa *et al*., 2022), its expression could be linked to a downstream stress signal rather than specifically nitrogen, while the concomitant *NIT1*/*NIT2* regulation may reflect the auxin response rather than nitrogen sensing per se. The most economical explanation is that biuret engages signalling pathways that normally monitor stress or defence-related cues, co-opting modules that couple perception to meristem activity. This hypothesis is consistent with our data without being proven by them: we identify neither a sensor nor a direct target, and ^15^N tracing cannot distinguish intact biuret from possible nitrogen-rich derivatives (Ochiai *et al*., 2022). In rice, excess biuret elicits oxidative-stress markers (elevated malondialdehyde) and a transcriptional response that predominantly down-regulates genes for chloroplast development and photosynthesis in leaves, rather than targeting cell division in roots (Zhang *et al*., 2023). This interspecific comparison, that must be taken carefully, indicates that biuret can mobilise stress responses in more than one species, although whether common signalling modules are shared across organs and species remains to be established.

Taken together, our results support an integrated working model (Figure 9) that links four demonstrated features of the response: *bir29* resists despite taking up and accumulating as much biuret as the wild type; biuret shifts auxin- and cytokinin-responsive outputs at the root tip; the transcriptional programme of *bir29* is reconfigured relative to the wild type; and local biuret exposure also inhibits the growth of untreated roots in split-root assays, revealing a systemic component. Several questions rise directly from this model: whether manipulating auxin or cytokinin signalling repositions the growth set-point; whether the stress- and defence-like signature reflects a functional response or remains correlative; and whether the systemic component reflects a generated long-distance signal or the translocation of biuret itself. Addressing them would clarify how a compound that land plants do not directly catabolise gains access to endogenous signalling networks, and whether a dedicated sensing mechanism is involved. Answering that would turn biuret from an agronomic nuisance into an informative probe of how root meristems translate chemical perturbations into developmental decisions.

**Figure 9.**
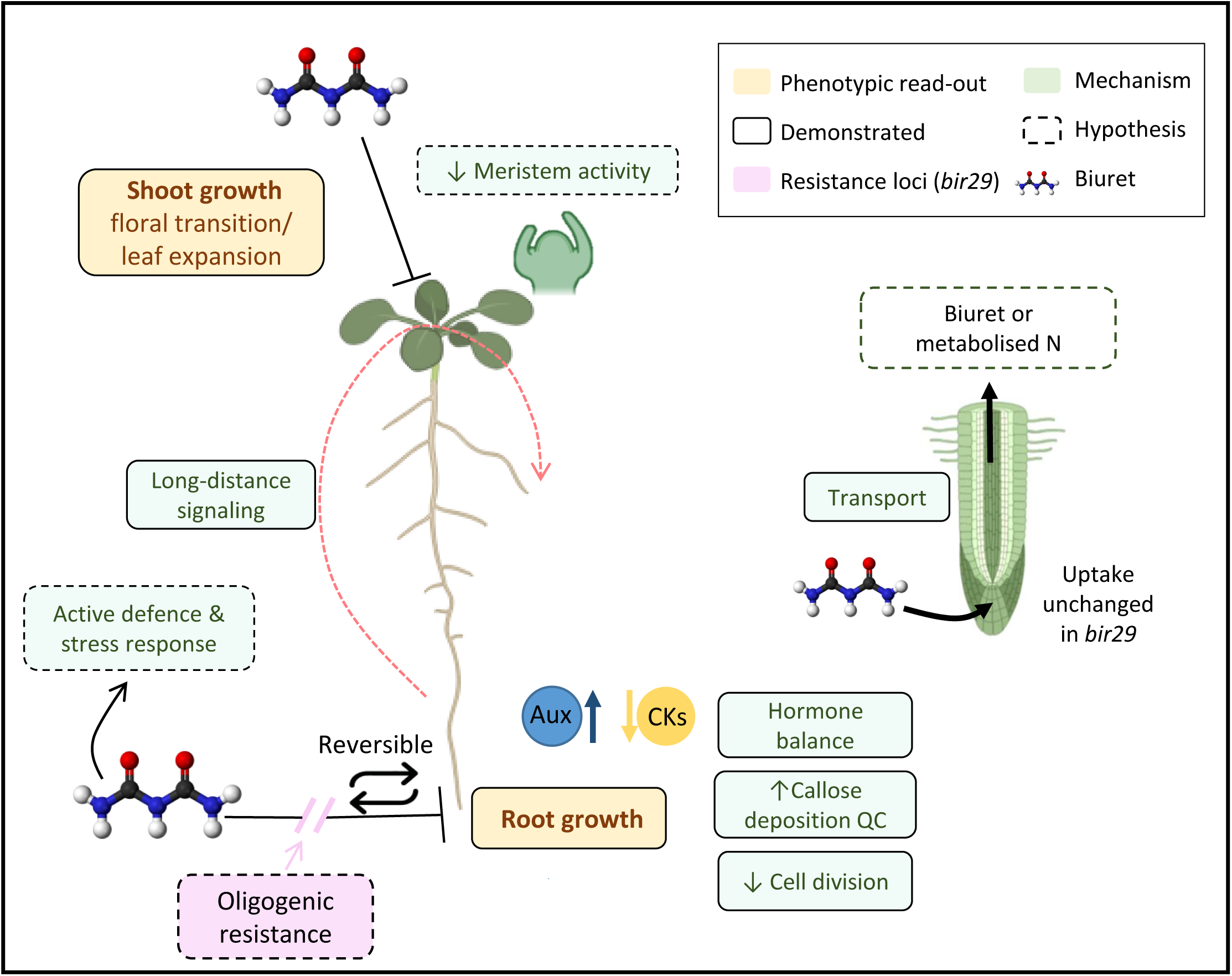
Integrated model of biuret effects on *Arabidopsis thaliana* development and the proposed mechanisms underlying the phenotype. Schematic summary of the demonstrated responses of *Arabidopsis* to biuret and of the working hypotheses derived from this study. Biuret inhibits both shoot and root growth (light yellow read-outs). In the shoot, foliar application reduces leaf expansion and impairs the floral transition. In the root, biuret triggers a dose-dependent inhibition of primary root growth that is fully reversible upon transfer to biuret-free medium (Reversible). At the cellular level (light green, solid borders), this inhibition is associated with a reduced number of meristematic cells and a decreased pCYCB1;1 reporter activity (↓ Cell division), with callose deposition around the quiescent centre (↑ Callose deposition QC), and with a reorganisation of hormonal outputs at the root tip, where the auxin-responsive DR5 signal increases (Aux, blue) and the cytokinin-responsive TCS signal decreases (CKs, yellow); because these reporters monitor transcriptional outputs rather than hormone concentrations, ‘Hormone balance’ refers to this shift in responsive activity. Biuret is taken up by the root and translocated within the plant (Transport, black arrows). Split-root experiments further show that, in addition to a predominant local effect, biuret elicits a systemic response that restrains growth in the untreated root system and is abolished in the resistant *bir29* mutant (Long-distance signaling). The biuret-resistant mutant *bir29* carries two T-DNA insertions, at *At1g48820* (*TPS*) and in the *PBL1*–*SCL30* intergenic region. Biuret uptake and accumulation are unchanged in *bir29* relative to the wild type, indicating that resistance is uncoupled from uptake and acts downstream, at the level at which biuret-induced perturbations are translated into the growth response. In the model, *bir29* therefore interrupts (/ /) the inhibitory route linking biuret perception to root growth arrest rather than reducing biuret entry. Hypotheses are shown with dashed borders. They comprise a reduction of shoot meristem activity (↓ Meristem activity); the engagement of an active defence and stress response (Active defence & stress response), inferred from the pathogen, reactive oxygen species and stress-related transcriptional signature induced in wild-type but not in *bir29* roots and not yet validated functionally; and the unresolved identity of the perceived and translocated species, that is whether it is biuret itself or a nitrogen-rich derivative, which total ¹⁵N quantification cannot distinguish (Biuret or metabolised N?). Box fill indicates the category of each element: phenotypic read-outs (light yellow) and mechanisms or molecular processes (light green). Box border indicates the level of supporting evidence: solid borders denote experimentally demonstrated features, whereas dashed borders denote hypotheses that remain to be tested functionally. The genetic determinant identified in this study is shown as a pink ellipse (*bir29*). Biuret is represented by its ball-and-stick structure; flat-ended bars (⊥) denote growth inhibition, black arrows denote transport, and the red dashed arrow denotes long-distance signalling.

## General conclusion

Biuret has long been viewed as a toxic by-product of urea fertilization (Mikkelsen, 1990), acting mainly through passive chemical injury (Achor and Albrigo, 2005). Our work reframes this view. In Arabidopsis, biuret does not merely poison growth: it elicits a structured (Figure 1), reversible (Figure 1f) and genetically tractable (Figure 3, 4) developmental response. This response engages major hormonal signaling outputs, including auxin and cytokinin pathways, propagates beyond the locally exposed tissue through a long-distance component (Figure 5), and can be genetically uncoupled from biuret uptake and accumulation (Figure 6). The identification of biuret-resistant genotypes whose phenotype cannot be explained by reduced exposure reveals that plants possess endogenous mechanisms that translate biuret perception into developmental decisions. Thus, biuret emerges not only as i) an agro-ecosystem contaminant, a ii) rhizosphere product of microbial activity, but also now as a potential signaling molecule. We believe that this shift in our understanding can lead to the development of genotypes being adapted to this molecule shown to be detrimental to plant adaptation and yield.

## Methods

### Plant material

*Arabidopsis thaliana* plants ecotype Columbia (Col-0) was used as wild type. The set containing 100 pools of 100 T-DNA insertion lines used for the genetic screening (N76508) were obtained from the NASC genetic ressources center. Transgenic lines used in this paper were pCYCB1;1::GUS (Colón-Carmona *et al*., 1999), pWOX5::GFP, DR5::GFP, DR5::GUS, TCS::GFP were kindly provided by colleagues. DR5::GFP (*bir29*) and TCS::GFP (*bir29*) lines were generated by crossing, and F2 progeny were screened to identify plants homozygous for the fluorescent reporter constructs and carrying the T-DNA insertions. Wild-type Col-0 and *bir29* crossing, followed by one self-pollination, obtained the F_2_ population.

### Plant growth conditions and treatments

For *in vitro* culture, seed were surface sterilized and cultivated in Petri dishes on MS without N (MS-N) media (PlantMedia) supplemented by 2.5mM KNO_3,_ sucrose 1% (w/v), MES 0.5% (w/v) and agar 10.5g.L^-1^ and cultivated vertically in a growth camber at 23°C, 120µmol photons m^-2^.s^-1^ and 16h photoperiod. For the root treatment condition, biuret stock solution (100mM) was filter sterilized and added to the media after sterilisation at the desired concentration. For shoot treatment condition, seeds were sawn on soil in pots covered with a plastic lid, and grown for three weeks in a growth chamber (16h day/night photoperiod, 23°C, 65% RH). Biuret was applied on leaves by spraying 1ml solution containing biuret 1% (w/v) and Silwet L-77 (0.05%) per rosette per day, during 14 days.

### Genetic screening

Six hundred seeds from each one of the 100 pools contained in the set N76508 obtained from the NASC were surface sterilized and sowed on MS-N supplemented by KNO_3_ 2.5mM and 1mM or 2mM biuret, stratified for 3 days at 4°C in the dark and grown vertically for 10 days in a growth chamber. Plates were then scanned and each individual plant showing a root length phenotype longer than the Col-0 used as control was selected and rescued on a media w/o biuret for 4 days. Plants were then transferred to the greenhouse for amplification and the collected seeds were phenotyped again on media containing 1mM or 2mM biuret as described previously.

### Split root experiment

For split root experiment, we adapted the protocol from (Ruffel *et al*., 2011). Seeds were surface sterilized, grown for 10 days in MS-N media containing 2.5mM KNO_3_ and after the cut of the primary root plants were transferred in split conditions were the compartments contained 2.5mM KNO_3_ with or without 2mM biuret (Biu2 and Biu0 respectively). Control plants (C) had roots in compartment separated but identical composition and Split plants (Sp) had roots in compartments with different composition in biuret. Seven days after transfer in split conditions, roots were scanned and PR length, LR number and LR density were measured by ImageJ.

### GUS staining

For DR5::GUS and pCYCB1;1::GUS, seeds were surface sterilized and sown in MS 2.5mM KNO3 with different concentrations of biuret solution. After 6 days seedlings were collected incubated in GUS staining buffer [0.1 M phosphate buffer pH 7, 0.1% (v/v) Triton X-100, 0.5 mM potassium ferricyanide, 0.5 mM potassium ferrocyanide, and 1 mM X-Gluc (5-bromo-4-chloro-3-indolyl-β-d-glucuronide)] and vacuum infiltration. The reaction was performed at 37 °C overnight in the dark after a 1 h vacuum treatment was applied at room temperature. After the reaction, samples were rinsed with distilled water and treated with 70% (v/v) to 10% (v/v) ethanol. Images were captured by using optical sectioning microscope Apotome (Zeiss).

### Root meristem imaging and cortical cell counting

The pWOX5::GFP plants, used as a marker of the quiescent center, were grown on control medium (MS-N supplemented with 2.5 mM KNO₃) or on control medium supplemented with 1 mM biuret. After 6 days of growth, root cell walls were stained with propidium iodine 10 mg.ml^-1^ (Sigma) and observed using a confocal laser scanning microscope (Zeiss LSM510). Root cells were counted starting from de pWOX5::GFP signal to the first elongating cell.

### Callose staining

Callose staining was performed on six-days old seedlings. Plantlets were collected and fixed for 10 minutes in the FEW solution (2% (v/v) formaldehyde, 63% (v/v) ethanol, 35% (v/v) water). After 1 minute incubation in sodium hydroxide 0.1M, plants were rinsed twice in phosphate buffer (0.1M K-Phosphate pH=7.2), then incubated 1.5 hours in aniline blue staining solution (0.1% w/v aniline blue in K-Phosphate buffer). Roots were observed using fluorescence microscope (excitation 390nm, emission 460 nm).

### Microscopy

DR5::GFP, TCS::GFP plants were cultivated for 6 days in control or biuret containing (0.5mM, 1mM or 2mM) media, using an optical sectioning microscope Apotome (Zeiss). Samples were excited at 470 nm and the GFP emission was collected at 525nm. GFP signal intensities were quantified using ImageJ.

### Biuret uptake

For the long term uptake experiment, wild-type and *bir29* plants were cultivated for three weeks in hydroponic system (media composition: KH_2_PO_4_ 1mM, du MgSO_4_,7H_2_O 1,5 mM, du K_2_SO_4_ 0,25 mM, du CaCl_2_ 3mM, du KNO_3_ 2.5mM et Na-Fe-EDTA 0,1 mM, H_3_BO_3_ 30μM, MnSO_4_,H_2_0 5μM, ZnSO_4_,7H_2_O 1μM, CuSO_4_,5H_2_0 1μM,(NH_4_)_6_Mo_7_O_24_ 0,1μM et Kl 5μM). ^15^N-biuret (Sigma) was added in the growth media at a final concentration of 0.5mM (10% atom excess ^15^N). After 3 days, plant were collected, rinsed in 0.1 mM CaSO_4_, and roots and shoots sparated prior to the analysis. For the influx rate determination, wild – type and *bir29* plants were cultivated in solid control media (see previous paragraphs) in vertical Petri dishes for 12 days. Then plants were transferred to 0.1 mM CaSO_4_ solution for 1 min, then to ^15^N-biuret 1mM solution (99% atom excess ^15^N) for 5 minutes and finally washed in CaSO_4_ solution for 1 min. Roots were then separated from shoots, and analysed using a Euro-EA Eurovector elemental analyzer coupled with an IsoPrime mass spectrometer(GV instruments, Crewe, UK), in the Isotopes Quantifications platform (AQUI, IPSiM, France).

### Transcriptomic analysis

For transcriptomic analysis, we grown wild-type and *bir29* mutant plants for 6 days in MS-N (+2.5mM KNO_3_) media supplemented with 0 or 1mM biuret. Root tissue samples were collected from 60 seedlings from control condition and 75 seedlings from biuret 1mM condition for each genotype. The experiment was repeated 3 times. Total RNA was extracted using the TRI Reagent™ method; 1µg was treated with DNAseI (Sigma) followed by a phenol:chloroform purification. RNA integrity was checked by microfluidic analysis in an Agilent 2100 Bioanalyzer. Following the manufacturer protocol, 100ng of total RNA were used to obtain 5.5µg of cDNA that were further fragmented, labelled and hybridized on the GeneChip ™ Arabidopsis Gene 1.1 ST Array Strip (Appliedbiosystems) and processed using the GeneAtlas®System. Data analysis was performed with R, a two way ANOVA model was applied followed by a Tukey *post-hoc* test. Differential expressed genes with ANOVA *p*<0.05 and Tukey adj. *p*ues <0.05 were considered for further analysis. Hierarchical clustering and Subset Intersections size analysis were performed with *Pheatmap* and *Vendetails* R packages respectively. GO terms analysis was performed using the https://geneontology.org/ web database.

### Whole-genome resequencing and identification of T-DNA insertions

Wild-type (Col-0) and *bir29* genomic DNA was isolated using the CTAB method (Doyle, 1987 #1630) and whole-genome sequencing libraries were prepared and sequenced by Novogene Co., Ltd. (Beijing, China) on an Illumina platform (e.g., NovaSeq X Plus), according to the manufacturer’s protocols. Raw sequencing data were mapped to the *Arabidopsis thaliana* TAIR10 reference genome by Novogene. Alignment files were visualized and manually inspected using Integrative Genomics Viewer (Robinson et al., 2011) with the TAIR10 genome assembly and *Arabidopsis_thaliana.TAIR10.59.gff3* annotation file.

### PCR genotyping and Sanger sequencing of insertion loci

Fragment amplification for genotyping was performed using the GoTaq® G2 Polymerase (Promega, Madison, Wisconsin, USA), following the manufacturer’s instruction, using the PCR primers listed in Table S9. The Sanger sequencing was performed by Eurofin Genomics (Ebersberg, Germany) on the amplified fragment isolated from gel after electrophoresis on 1% (w/v) agarose gel separation or directly from the PCR reaction, using the same primers of the amplification reaction.

### Statistical analysis

Experiment were performed at least three times. Statistical significance of genotype, treatment or genotype*treatment factors was assessed by parametric or non-parametric test, based on data normality and homoscedasticity, as indicated in the legends and applying a significance threshold of 0.05. Statistical analysis was performed using RStudio (Posit, Boston, MA, USA) or GraphPad Prism 10.0.1 (Graph-Pad Software, San Diego, CA, USA).

## Data statement

All raw and processed data generated in this study have been deposited in the EBI data base. The genomic resequencing data for Col-0 and *bir29* mutant are available in the ArrayExpress database (http://www.ebi.ac.uk/arrayexpress) under the accession number E-MTAB-17253. Microarray data are available under accession number E-MTAB-17283.

## Disclosure of AI Use

During the preparation of this manuscript, the authors used ChatGPT to assist with the language editing of portions of the text. The tool was not used to design the experiments, nor to generate, collect, analyse or interpret the experimental data, and it was not used to create, alter or manipulate any figures. After using this tool, the authors reviewed and edited all content as needed and take full responsibility for the accuracy and integrity of the work and for the final version of the manuscript.

## Conflict of Interest statement

The authors declare no conflict of interest.

## Acknowledgments

This work was supported by the Agence Nationale de la Recherche (ANR; ANR- ANR-21-CE20-0008 – SNUpReg; to A.M.) and Institut Agro Montpellier (CRI-U-LEAF to A.M.). We acknowledge Cécile Fizames for assistance with genomic data analyis, the Isotopes Quantifications Platform (AQUI, IPSIM, France) for isotope quantification, and the Montpellier Ressources Imagerie (MRI) platform for assistance with image acquisition and quantification.

## Author contributions

VP performed experiments and the data analysis, and wrote the manuscript, VT, AD and TP contributed to performing experiments and data analysis, BL and GK revised the manuscript, AM designed the research, performed data analysis and wrote the manuscript.

## Notes

http://www.ebi.ac.uk/arrayexpress

### Competing Interest Statement

The authors have declared no competing interest.

