## Supplementary material for "Biuret inhibits Arabidopsis root growth through an active, reversible, and genetically tractable developmental response": Figure S1-S8

### Fig S1

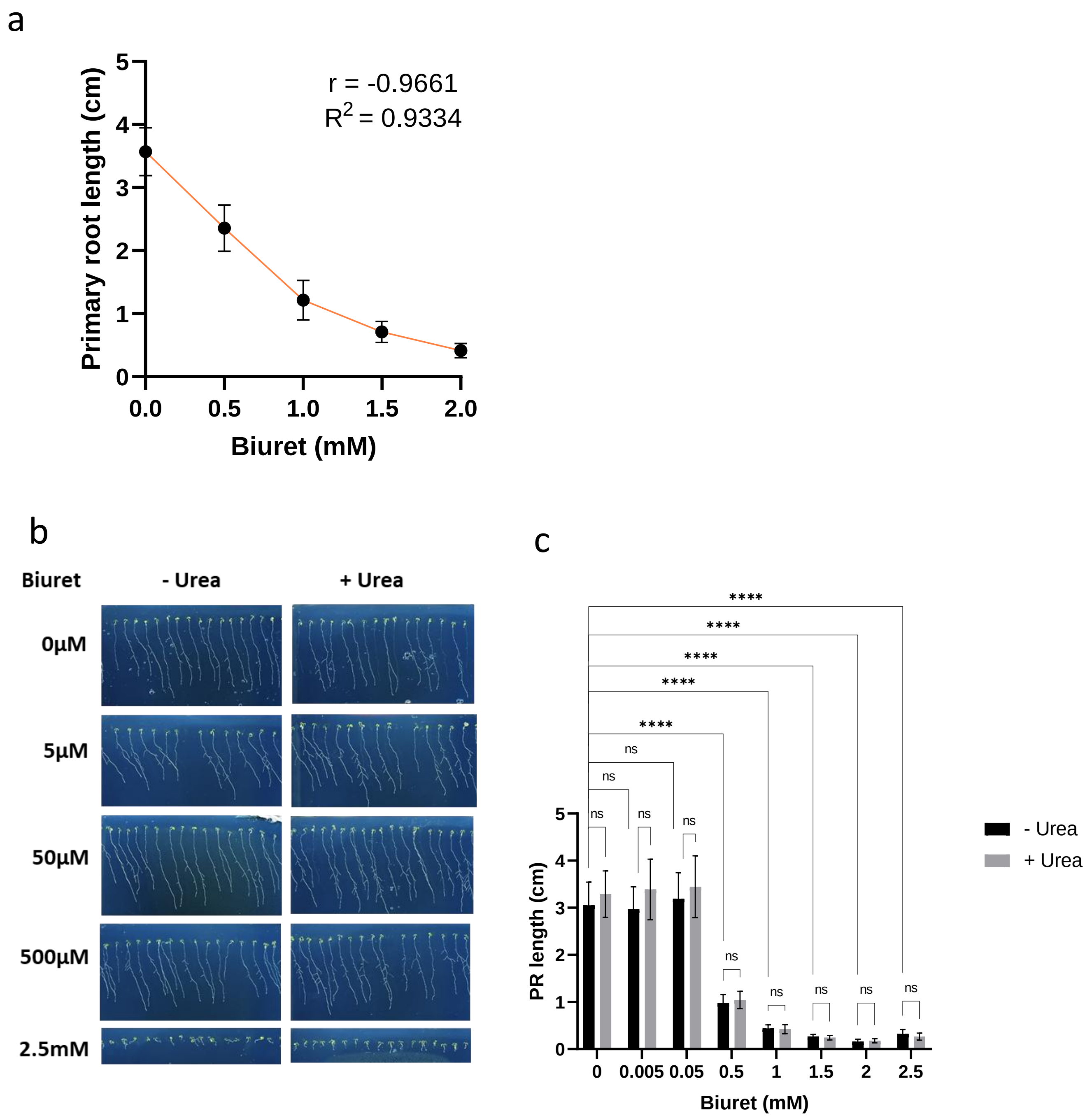


**Fig. S1. Primary root growth dose response to high and low biuret concentrations and interaction with urea.**
(a) Primary root growth dose response to biuret; Pearson r coefficient and $R^2$ for the correlation analysis are indicated. (b) Representative phenotypes of seedlings grown for 13 days on media containing increasing concentrations of biuret (5–500 µM) in the absence (−urea) or presence (+urea) of 2.5 mM urea. (c) Quantification of primary root length after 13 days of growth under the same conditions. Bar plts represent mean±SD. Asterisks indicate significant differences relative to the corresponding 0 µM biuret condition. ns indicates no significant difference. (*****p* < 0.0001; one-way ANOVA followed by Tukey's post hoc test).

### Fig.S2

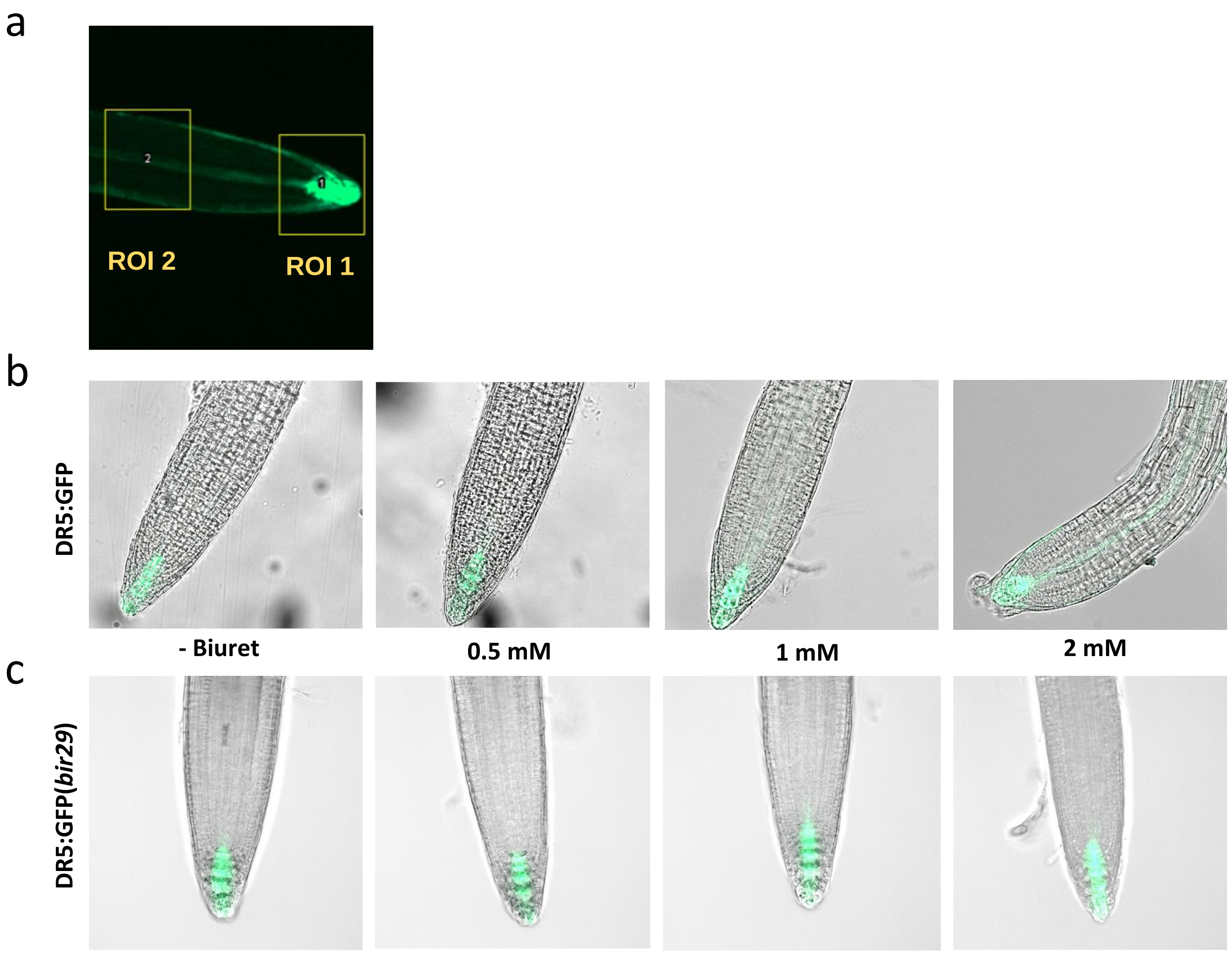


**Fig. S2. Biuret enhances auxin-responsive DR5:GFP activity at the root tip.**
(a) Region of interest (ROI) defined along the primary root and used for DR5:GFP fluorescence quantification in Fig. 2.
(b,c) Representative epifluorescence images of DR5:GFP seedlings of WT (b) or *bir29* genetic background mutant (c) grown for 11 days on media containing 0, 0.5, 1 or 2 mM biuret.

### Fig.S3

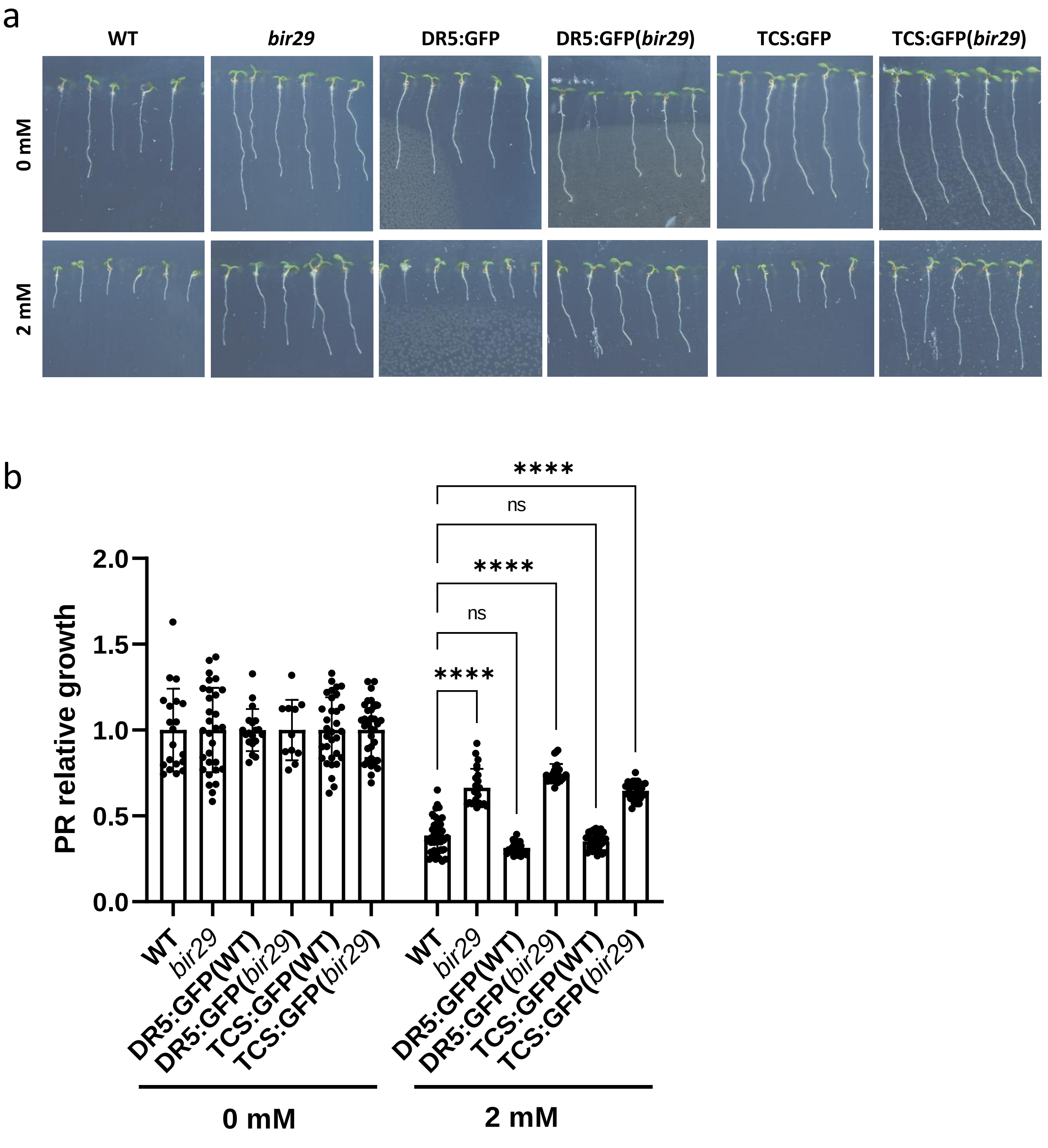


**Fig. S3. The bir29 mutation confers biuret resistance in DR5:GFP and TCS:GFP reporter backgrounds.**

(a) Representative primary root phenotypes of WT, *bir29*, DR5::GFP, DR5::GFP(bir29), TCS::GFP and TCS:GFP(bir29) seedlings grown for 6 days on media containing 0 or 2 mM biuret. (b) Quantification of relative primary root growth in the different genotypes. For each line, primary root length values were normalized to the mean value of the corresponding 0 mM condition (n = 12–24). Asterisks represent significant differences between WT and *bir29* genotypes (****$p$ < 0.0001; two-way ANOVA followed by Dunnett's post hoc test).

### Fig.S4

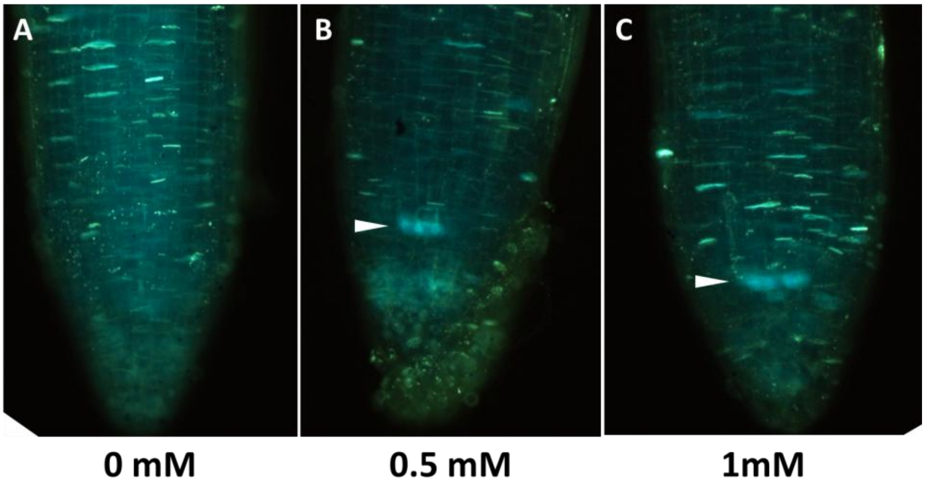


**Fig. S4. Biuret induces callose deposition at the root apex.**

(a) Representative fluorescence images of callose staining in roots of WT seedlings grown for 6 days on media containing 0, 0.5, 1 or 2 mM biuret. White arrows indicate callose deposits in the quiescent center (QC) region.

### Fig. S5

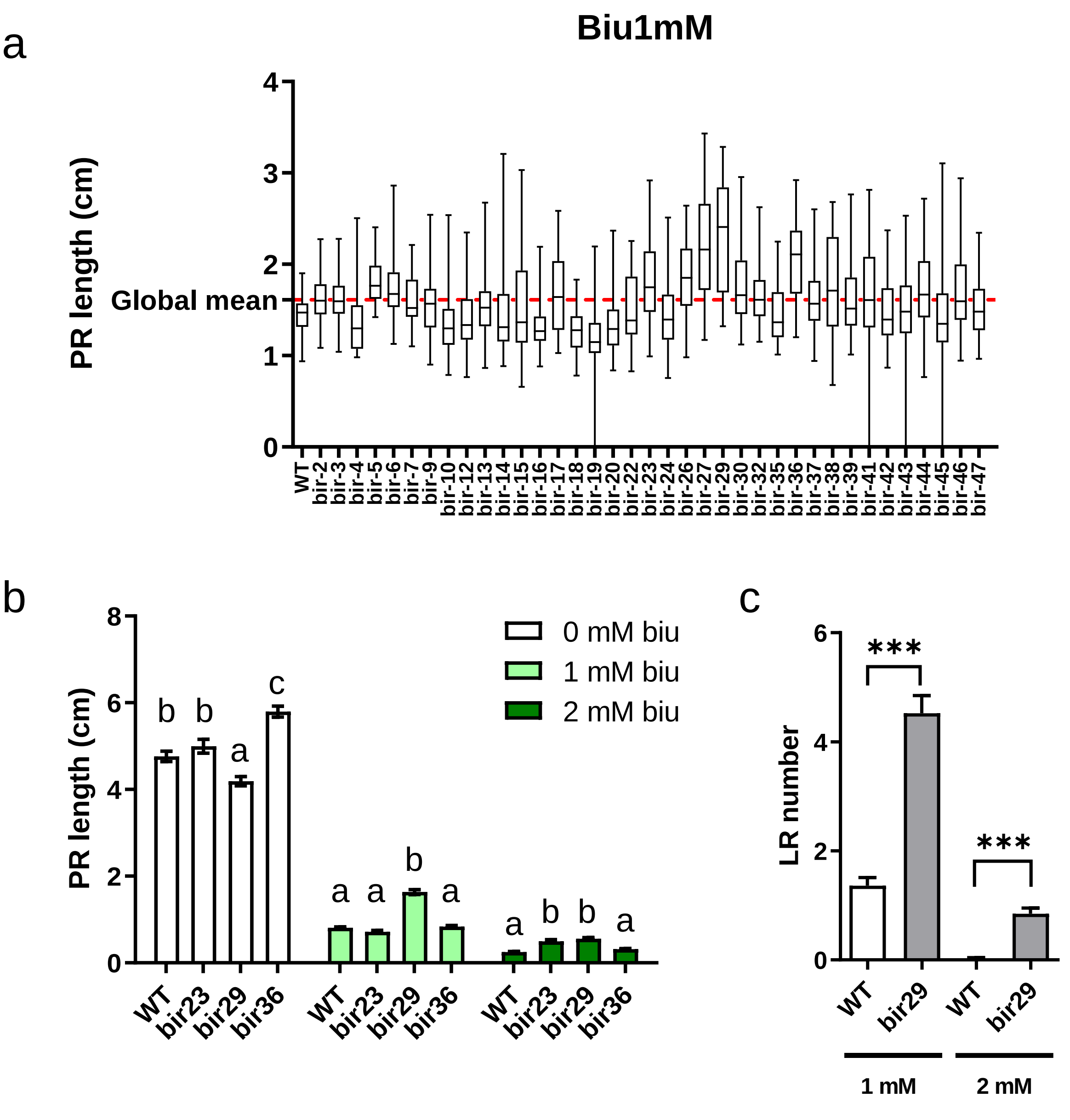


**Fig. S5. Screening of a T-DNA insertion population for altered sensitivity to biuret.**
(a) Phenotypic screening of the SALK T-DNA insertion collection for biuret sensitivity. The 37 pre-selected mutant lines were grown for 13 days in vitro on half-strength MS medium containing 1 mM biuret. Primary root length (PR) was measured for 30–70 plants per line. The dashed red line indicates the global mean PR length across all screened lines. (b) Primary root phenotype of the three selected lines (bir23, bir29 and bir36) displaying the largest difference in PR length relative to WT (Col-0) (n = 70–90). Plants were grown for 13 days in vitro on half-strength MS medium containing 0, 1 or 2 mM biuret. Different letters indicate groups with significantly different means ($p$ < 0.05; ANOVA followed by Fisher's LSD post hoc test). (c) Lateral root number in WT and bir29 plants grown on media containing 1 or 2 mM biuret. Asterisks indicate significant differences (***$p$ < 0.01; Student's t-test).

### Fig. S6

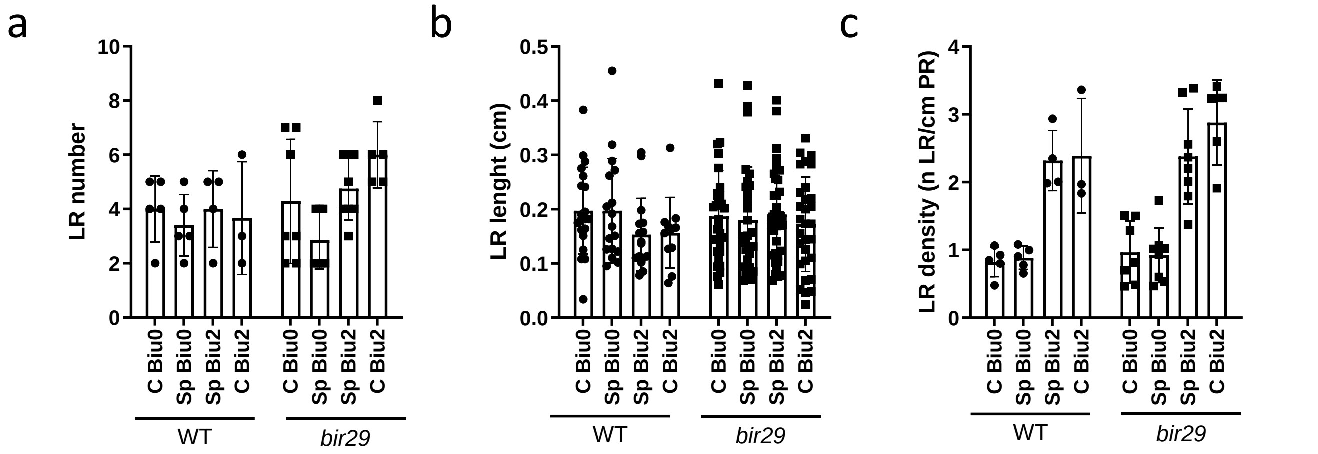


**Fig. S6. Local and systemic effects of biuret on root development in the *bir29* mutant.** (a–c) Quantification of lateral root number (a), lateral root length (b) and lateral root density (c) in wild type and *bir29* plants grown in a split-root system. Seedlings were initially grown on basal medium and subsequently transferred to the split-root system for 6 days. C Biu0 and Sp Biu0 correspond to biuret-free conditions, whereas Sp Biu2 and C Biu2 contain 2 mM biuret. In control plants (C), both root halves were exposed to the same medium, whereas in split plants (Sp), the two root halves were exposed to different media. Asterisks indicate significant differences ($p < 0.05$; one-way ANOVA followed by Tukey's post hoc test).

### Fig. S7

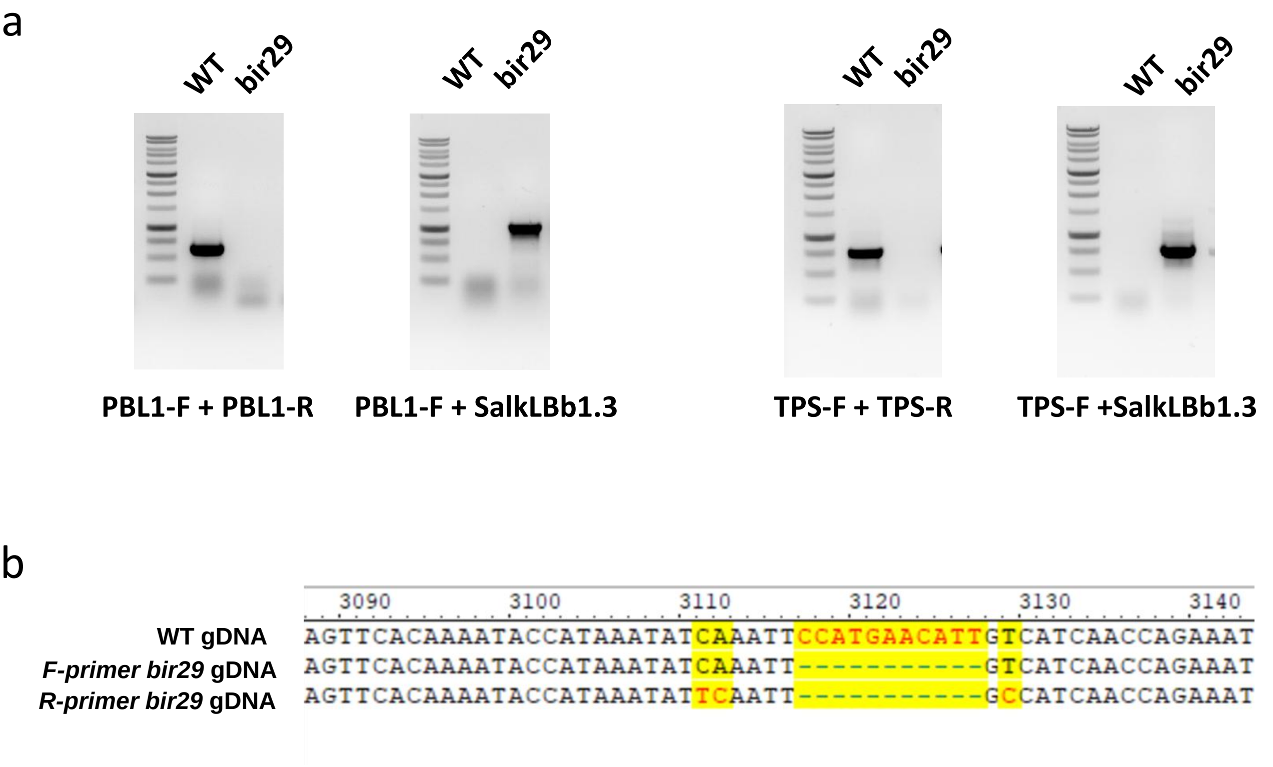


**Fig. S7. Identification of the T-DNA insertion site in the bir29 mutant.**
(a) Genotyping of *bir29* loci containing T-DNA insertion. The wild type and mutant regions corresponding to the intergenic insertion were amplified using PBL1-F and PBL1-R primer and the SalkLBb1.3 specific primers; the wild type and mutant regions corresponding to the insertion in TPS were amplified using TPS-F, TPS-R, and SalkLBb1.3 specific primers. (b) Sequence analysis of the *bir29* genomic DNA revealing a T-DNA insertion in the intergenic region between At3g55450 and At3g55460, together with an 11-bp deletion (highlighted in yellow) located 142 bp downstream of the 3′ UTR of the At3g55450.1 (PBL1) transcript. Sequencing was performed using a gene-specific forward primer (F-primer) and a T-DNA-specific reverse primer (R-primer).

### Fig. S8

a

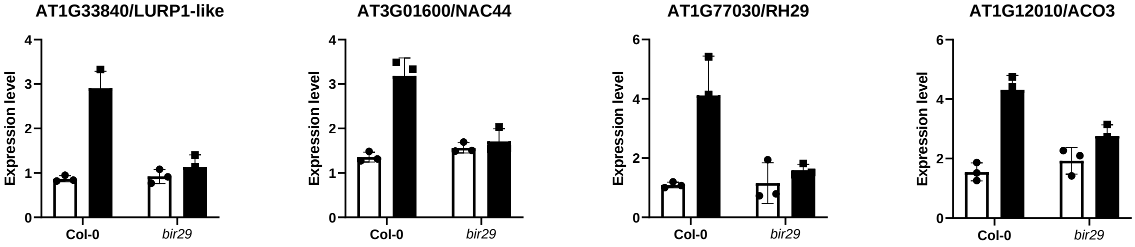


b

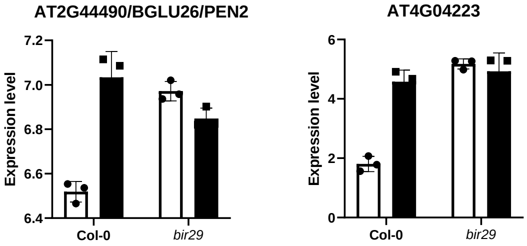


c

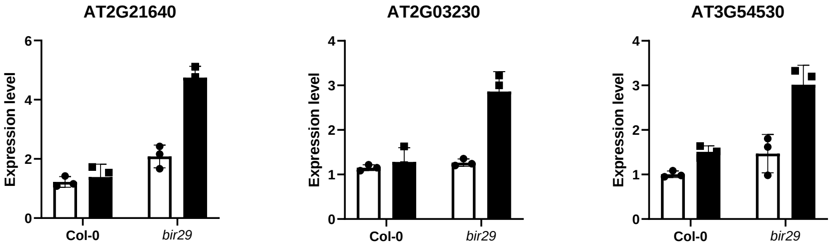


**Fig. S8. Transcriptional regulation of some interesting genes showing specific response in WT and bir29 plants.**

Transcriptomic expression levels of selected differentially expressed genes (DEGs) from the transcriptome dataset (Fig. 7), grouped by profiles: (a) upregulates in wild-type and constitutive low expression level in *bir29*; (b) upregulated in wild-type and constitutive high expression level in *bir29*; and (c) constitutive expression level in wild-type and upregulated in *bir29* cell-cycle regulation and root growth. For each gene, expression values are shown in WT (Col-0) and *bir29* plants under control (white bar) and biuret-treated conditions (black bar). Bars represent the mean±SD (n=3) and individual data points are overlaid.
