## Supplementary material for "Biuret inhibits Arabidopsis root growth through an active, reversible, and genetically tractable developmental response": Table S9

**Table S8.**

**Primers used for Genotyping and Sanger sequencing**

| Name | 5’-3’ Sequence |
| --- | --- |
| PBL1-F | GACCAAGTGGTTCGAGCCTT |
| PBL1-R | TTCCATGAACATTGTCATCAACCAG |
| TPS-F | GACAGCCAATTGAATAAGAAAAC |
| TPS-R | TATCTCTCTCTCAAGCGCATC |
| SalkLBb1.3 | ATTTTGCCGATTTCGGAAC |
